# Biological and genomic resources for the cosmopolitan phytoplankton *Bathycoccus:* Insights into genetic diversity and function of outlier chromosomes

**DOI:** 10.1101/2023.10.16.562038

**Authors:** Louis Dennu, Martine Devic, Janaina Rigonato, Angela Falciatore, Jean-Claude Lozano, Valérie Vergé, Cédric Mariac, Nathalie Joli, Olivier Jaillon, François Sabot, François-Yves Bouget

## Abstract

1.

Population-scale genome sequencing has become essential for exploring genetic diversity and adaptation, particularly in land plants. In contrast, eukaryotic phytoplankton resources remain limited to model reference genomes or community-level metagenomics, leaving a gap in understanding intraspecific variation and evolutionary processes. To address this, we developed a comprehensive biological and genomic resource for the cosmopolitan and ecologically important genus *Bathycoccus*. Extensive metagenomic data from across the world Ocean are available for this genus, and previous studies have identified four *Bathycoccus* species and reconstructed 34 metagenome-assembled genomes. Here we report 28 high-quality strain genome sequences using a combination of Oxford Nanopore Technologies long reads and Illumina short reads and associated biological resources. These include 24 *Bathycoccus prasinos* strains spanning a latitudinal gradient from 40° to 78°N, a reference genome for *Bathycoccus calidus*, and three genomes of the recently identified B3 clade, which we propose as the *Bathycoccus catiminus* species. Comparative analyses of sequenced genomes with MAGs highlight the complementarity between resources: while MAGs capture environmental diversity and uncover uncultured taxa, the cultured strain genomes provide complete, non-chimeric high-quality assemblies that resolve structural variations and haplotype-level diversity not detected in MAGs. These include the large outlier chromosome (BOC), a putative sexual chromosome revealing a second mating type, and extensive variability in the small outlier chromosome (SOC), associated with viral resistance and genome plasticity. Together, these biological and genomic resources establish *Bathycoccus prasinos* as a powerful model for studying diversity, adaptation, and evolution of eukaryotic phytoplankton in the ocean, complementing existing global metagenomic datasets.

**Significance statement:** Eukaryotic phytoplankton are key to ocean ecosystems, yet their intraspecific genomic diversity is poorly understood. We present 28 high-quality genomes and their annotations of the cosmopolitan microalga *Bathycoccus*, revealing whole genome structural variations, chromosomal haplotype diversity linked to mating and viral resistance, and offering a genomic framework that complements metagenomic data to establish this picoalga as a model for functional and ecological studies.

## 3. Introduction

Accessing the natural diversity of a species constitutes an invaluable resource, providing insights into its evolutionary history and facilitating correlations between phenotypes and genotypes through functional studies. In plants, the availability of extensive collections of natural accessions and crop cultivars from diverse environments has greatly improved our knowledge of adaptation (Alonso-Blanco et al., 2009). Furthermore, large-scale sequencing projects on widely distributed models, such as for *Arabidopsis thaliana*, and in several model crops, have provided a comprehensive framework for the study of genomic diversity, its underlying mechanisms and their impact on environmental adaptation (100 Tomato Genome Sequencing Consortium, 2014; 3,000 Rice Genomes Project, 2014; Alonso-Blanco et al., 2016). Other reference organisms have also emerged as models in population genomics, such as *Saccharomyces cerevisiae* for which extensive sequencing of available strains highlighted complex genomic structures with great diversity (Peter et al., 2018). While these comprehensive sequencing projects have benefited their research communities, the study of eukaryotic phytoplankton lags behind in terms of comparable large-scale sequencing initiatives on single species. Pioneer studies have been conducted for the marine diatom *Phaeodactylum tricornutum* or the freshwater and estuarine haptophyte *Prymnesium parvum* (Rastogi et al. 2020, Wisecaver et al., 2023). However, in most currently available pangenomic studies of eukaryotic phytoplankton, descriptions of intraspecific diversity are limited by the presence of cryptic species within taxa that were initially separated only as morphospecies, as exemplified for *Emiliana huxleyi* (*Gephyrocapsa huxleyi*) (Read et al., 2013; Bendif et al., 2023), *Microcystis* (Cai et al., 2023) or *Prymnesium parvum* (Wisecaver et al., 2023). This highlights current challenges in the study of phytoplankton intraspecific diversity, as the taxonomic resolution and large morphological diversity of many phytoplankton taxa must be taken into account when characterizing intraspecific subpopulations and distinguishing cryptic species that have undergone speciation events. Although access to intraspecific diversity in eukaryotic phytoplankton is constrained by the limited availability of biological resources for most species, the rise of large metagenomic datasets over the past 15 years has enabled the reconstruction of metagenome-assembled genomes (MAGs), providing a complementary resource for exploring diversity (Delmont et al., 2022, Duncan et al., 2022, Xu et al., 2024).

The global distribution and high abundance of phytoplankton species make them major actors of the planet’s primary production, contributing as much as land plants (Li, 1994; Worden et al., 2004). In particular, eukaryotic phytoplankton species are characterized by their great taxonomic diversity and relatively short generation times allowing for their ecological success in a wide range of environmental conditions in the world Ocean (Lynch et al., 1991; De Vargas et al., 2015). Microalgae also serve as valuable models for both fundamental studies and biotechnological research. Among eukaryotic phytoplankton, the *Mamiellophyceae* class stands out as unicellulars widely distributed in the oceans from poles to equator and marked by seasonal population dynamics in polar and temperate regions (Lambert et al., 2019; Leconte et al., 2020). *Mamiellophyceae* diverged at the base of the green lineage, making them attractive models for studying essential cellular functions from an evolutionary perspective (Leliaert et al., 2012; Yung et al., 2022). The early whole genome sequencing of *Ostreococcus tauri* and the implementation of genetic transformation tools, including gene targeting by homologous recombination, have established *O. tauri* as a model for the study of several cellular pathways such as the circadian clock, the cell division, iron metabolism and vitamin B1 metabolism (Derelle et al., 2006; Corellou et al., 2009; Moulager et al., 2010; O’Neill et al., 2011; Van Oijen et al., 2013; Lozano et al., 2014; Botebol et al., 2015; Paerl et al., 2017; De Barros Dantas et al., 2023). The genomic content of *O. tauri* unveiled several features shared by *Mamiellophyceae*: A compact haploid genome, a majority of monoexonic genes, limited gene redundancy and two outlier chromosomes, characterized by lower GC content and unique gene structures, named big outlier chromosome (BOC) and small outlier chromosome (SOC), respectively (Derelle et al., 2006; Grimsley et al., 2015). The BOC is putatively involved in mating mechanisms, as evidenced by the identification of two haplotypes in several *Mamiellophyceae*, and potential recombination suppression, but experimental evidence for sexual reproduction in *Mamiellophyceae* is still lacking (Blanc-Mathieu et al., 2017). The SOC is hypervariable between strains and has been shown to be associated with viral resistance mechanisms (Yau et al., 2016; Blanc-Mathieu et al., 2017). While *O. tauri* offers valuable insight into the origin and evolution of biological processes in the green lineage, its low abundance in marine environments, as inferred from metagenomic dataset, hinder its broader use in investigating questions relevant to ecology and adaptation (Demir-Hilton et al., 2011).

In addition to *Ostreococcus*, the *Bathycoccaceae* family also includes the *Bathycoccus* genus, a picoalga characterized by its scales-covered cell (Eikrem & Throndsen, 1990). Mostly studied through metabarcoding and metagenomic approaches, there are four *Bathycoccus* species, initially defined as a single species *Bathycoccus prasinos*, but later split into distinct evolutionary clades (B1, B2, B3 and B4) (Moreau et al., 2012; Bachy et al., 2021; Xu et al., 2024). *B. prasinos* (B1) *per se* shows high abundance at high latitudes between temperate and polar regions, as opposed to *B. calidus* (B2), mostly found in warm oligotrophic waters at lower latitudes (De Vargas et al. 2015; Vannier et al., 2016; Leconte et al., 2020). For the recently characterized B3 and B4 clades, B3 is particularly abundant in the China Sea, while B4 occurs primarily in the Baltic Sea. Notably, a representative strain of the B3 clade was isolated from the Bay of Hong Kong and sequenced, providing additional genomic resources for this lineage (Xu et al., 2024).

Among *Bathycoccus* species, *B. prasinos* displays the widest geographic distribution along latitudinal gradients and exhibits strong seasonal patterns in both temperate and arctic waters, with population growth often occurring through annual bloom events (Joli et al., 2017; Lambert et al., 2019; Devic et al., 2024). The cosmopolitan distribution of *B. prasinos* from polar environment, marked by dramatic changes in photoperiod and temperature, to temperate Mediterranean climate makes this phytoplankton a prime model for the study of both latitudinal and seasonal adaptation mechanisms. The reference genome of *B. prasinos* is of 15 Mb in size and is composed of 19 nuclear chromosomes supporting 7,847 annotated genes (Moreau et al., 2012). It features, as for other *Mamiellophyceae*, a BOC and a SOC, corresponding to chromosomes 14 and 19, respectively. Additional genomic resources comprise single-cell assembled genomes of both *Bathycoccus* species and several environmental metagenome-assembled genomes, but the variable completion rate of genomes produced by these methods currently provides a fragmented and incomplete view of the genetic diversity within the *Bathycoccus* genus (Vaulot et al., 2012; Vannier et al., 2016; Joli et al., 2017; Benites et al., 2019; Delmont et al., 2022).

In this study, we aimed to draw a comprehensive landscape of *Bathycoccus* genetic diversity at the intraspecific (*i.e* within *B. prasinos*) levels and to provide a biological and genomic resource for future studies of molecular mechanisms underlying adaptation to environmental niches in *B. prasinos*. For this purpose we selected and sequenced a panel of *Bathycoccus* strains including (i) collection strains previously isolated in different geographic locations in north western europe, (ii) strains that have been isolated during the winter bloom of 2018-2019 in the Banyuls bay (Mediterranean sea, France) (Devic et al., 2024) and (iii) arctic strains from the Baffin bay (Arctic Ocean) we isolated and genotyped. Both ONT long reads and illumina short reads were used to generate 28 *de novo* genome assemblies of high quality and completeness. This resource was used to describe the genetic diversity of *Bathycoccus* at both species and intraspecific levels with a focus on outlier chromosomes and their putative functions.

## 4. Results

### Biological resource

The biological resource used in this study includes 256 strains of *Bathycoccus sp*. from different geographic locations including the Mediterranean Sea, English Channel, North Sea, Arctic Ocean and Indian Ocean. These comprise seven strains of *B. prasinos* selected from the Roscoff Culture collection (RCC5417, RCC1613, RCC685, RCC1615, RCC1868, RCC4222, RCC4752) and initially sequenced using ONT sequencing to identify polymorphic indel markers (Devic et al., 2024). These markers were used to characterize the genetic diversity of 66 *Bathycoccus sp.* strains isolated in the bay of Banyuls-sur-mer (Mediterranean sea, France) during the 2018/2019 winter bloom, allowing the selection of seven representative *B. prasinos* isolates for sequencing (G11, C2, G2, E2, A8, B8, A1) (Devic et al., 2024). Similarly, seawater and ice samplings were conducted in three stations of the Baffin bay as part of the DarkEdge cruise in October 2021. After incubation of microbial communities (sizes < 1.2 µm) at 4°C or 15°C, picophytoplankton clones were obtained in semi-solid low-melting agarose. Genotyping using *B. prasinos* specific primers of the TOC1 gene revealed that all 183 isolates corresponded to *B. prasinos*. Further genotyping using two polymorphic indel markers led to the identification of 10 multi-loci genotypes representative of the *B. prasinos* diversity in the Baffin bay. One isolate of each genotype was retained for sequencing (A818, B218, B518, C218, E318, H718, D119, H44, A727, A827) **(Table S1)**.

In total, we selected 24 strains of *B. prasinos* for sequencing, with origins covering a latitudinal gradient from 40°N to 78°N (**Table 1**). The only strain of the recently described species *Bathycoccus calidus* (RCC716) isolated from the Indian ocean (Bachy et al., 2021) was also included. However, 11 isolates from the Banyuls bay showed weak amplifications of the LOV histidine-kinase marker. Sequencing of the 18S rDNA confirmed their identity as *Bathycoccus sp.*, but variations in the ITS2 region revealed that they were phylogenetically related to the recently described *Bathycoccus* B3 clade, that is distinct from *B. prasinos* (B1), *B. calidus* (B2) and from *Bathycoccus* clade B4 (Xu et al., 2024) corresponding to ITS2 environmental sequences recovered from the Kara Sea (Belevich et al., 2021) **(Figure S1)**. Three of these B3 isolates were thus selected for sequencing (C3, G5 and G8).

**Table 1:** Metadata of sequenced strains.

|  | Species | Latitude<br>(North) | Longitude<br>(East) | Depth<br>(m) | Sampling site | Sampling date | Sample source |
| --- | --- | --- | --- | --- | --- | --- | --- |
| **ARCTIC OCEAN** |  |  |  |  |  |  |  |
| A818 | *B. prasinos* | 78.12 | -74.36 | 0 | Baffin bay DE310 † | 16/10/2021 | DarkEdge |
| B218 | *B. prasinos* | 78.12 | -74.36 | 0 | Baffin bay DE310 † | 16/10/2021 | DarkEdge |
| B518 | *B. prasinos* | 78.12 | -74.36 | 0 | Baffin bay DE310 † | 16/10/2021 | DarkEdge |
| C218 | *B. prasinos* | 78.12 | -74.36 | 0 | Baffin bay DE310 † | 16/10/2021 | DarkEdge |
| E318 | *B. prasinos* | 78.12 | -74.36 | 0 | Baffin bay DE310 † | 16/10/2021 | DarkEdge |
| H718 | *B. prasinos* | 78.12 | -74.36 | 0 | Baffin bay DE310 † | 16/10/2021 | DarkEdge |
| D119 | *B. prasinos* | 78.12 | -74.36 | 0 | Baffin bay DE310 † | 16/10/2021 | DarkEdge |
| H44 | *B. prasinos* | 76.03 | -77.23 | 0 | Baffin bay DE110 | 12/10/2021 | DarkEdge |
| A727 | *B. prasinos* | 75.95 | -85.58 | 20 | Baffin bay DE410 | 21/10/2021 | DarkEdge |
| A827 | *B. prasinos* | 75.95 | -85.58 | 20 | Baffin bay DE410 | 21/10/2021 | DarkEdge |
| RCC5417 | *B. prasinos* | 67.48 | -63.78 | 0 | Baffin Island | 01/06/2016 | RCC ‡ |
| **ATLANTIC OCEAN** |  |  |  |  |  |  |  |
| RCC1613 | *B. prasinos* | 57.57 | 8.67 | 35 | North Sea | 26/07/2007 | RCC ‡ |
| RCC685 | *B. prasinos* | 54.18 | 7.90 | 0 | North Sea | 16/05/2001 | RCC ‡ |
| RCC1615 | *B. prasinos* | 50.2 | 0.32 | 10 | English Channel | 08/07/2007 | RCC ‡ |
| RCC1868 | *B. prasinos* | 48.75 | -3.95 | 0 | English Channel | 05/02/2009 | RCC ‡ |
| **MEDITERRANEAN SEA** |  |  |  |  |  |  |  |
| RCC4222 | *B. prasinos* | 42.48 | 3.53 | 3 | Banyuls bay | 01/01/2006 | RCC ‡ |
| G11 | *B. prasinos* | 42.48 | 3.53 | 3 | Banyuls bay | 03/12/2018 | OOB § |
| C2 | *B. prasinos* | 42.48 | 3.53 | 3 | Banyuls bay | 28/01/2019 | OOB § |
| G2 | *B. prasinos* | 42.48 | 3.53 | 3 | Banyuls bay | 11/02/2019 | OOB § |
| E2 | *B. prasinos* | 42.48 | 3.53 | 3 | Banyuls bay | 25/02/2019 | OOB § |
| A8 | *B. prasinos* | 42.48 | 3.53 | 3 | Banyuls bay | 26/02/2019 | OOB § |
| B8 | *B. prasinos* | 42.48 | 3.53 | 3 | Banyuls bay | 26/02/2019 | OOB § |
| A1 | *B. prasinos* | 42.48 | 3.53 | 3 | Banyuls bay | 19/03/2019 | OOB § |
| G8 | *Bathycoccus* sp. | 42.48 | 3.53 | 3 | Banyuls bay | 03/12/2018 | OOB § |
| C3 | *Bathycoccus* sp. | 42.48 | 3.53 | 3 | Banyuls bay | 12/12/2018 | OOB § |
| G5 | *Bathycoccus* sp. | 42.48 | 3.53 | 3 | Banyuls bay | 12/12/2018 | OOB § |
| RCC4752 | *B. prasinos* | 40.48 | 14.14 | 100 | Gulf of Naples | 17/04/1986 | RCC ‡ |
| **INDIAN OCEAN** |  |  |  |  |  |  |  |
| RCC716 | *B. calidus* | -14.48 | 113.45 | 70 | Indian Ocean | 11/06/2003 | RCC ‡ |

† Ice station
‡ Roscoff Culture Collection
§ Oceanological Observatory of Banyuls-sur-Mer

### Genomic resources

The 28 *Bathycoccus sp.* strains, corresponding to 24 *B. prasinos*, 1 *B. calidus* and 3 *Bathycoccus* B3 clade, display a wide geographical distribution ranging from arctic to equatorial regions and seasonal patterns in the Banyuls bay (Devic et al., 2024) (**Figure 1A**, **Table 1**). These strains were sequenced using both ONT and Illumina to produce long and short high quality reads respectively, ensuring contiguity and low error rates in the final assemblies. Genome assembly was conducted using a pipeline outlined in **Figure 1B**. ONT reads of approximative depth ranging from 12 to 300X, assuming a genome size of 15Mb (Moreau et al., 2012), were used for a genome assembly using FLYE and a post-assembly correction using Medaka. Illumina reads (30 to 130X depth) were used for final assembly polishing using Pilon. Scaffolding and orientation of the created contigs upon the reference genome from the strain RCC1105 (Moreau et al., 2012) was performed through RagTag.

**Fig 1:**
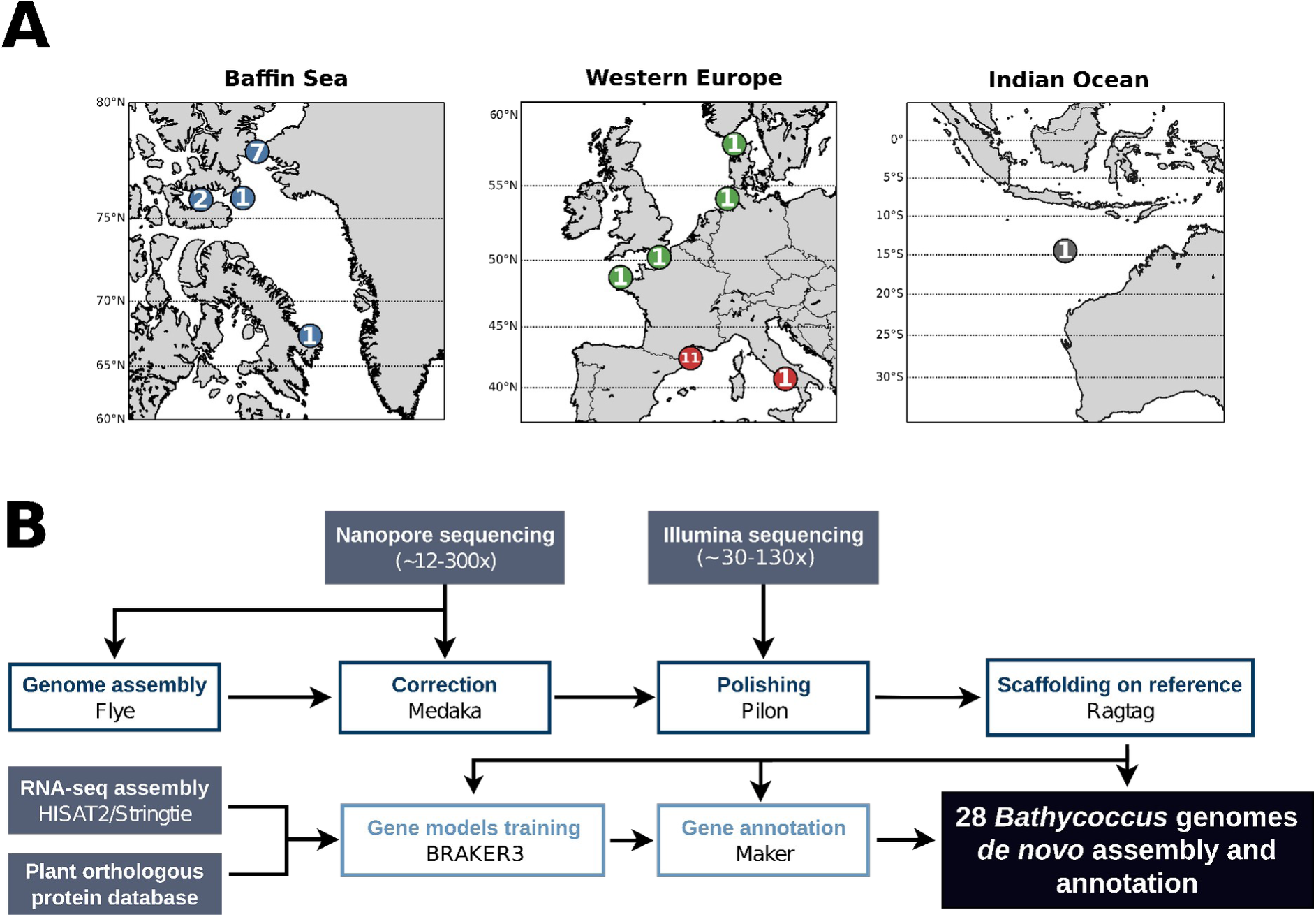
Sampling, sequencing, genome assembly and annotation strategy of *Bathycoccus*. (A) Geographical distribution of the sampling sites for the selected *Bathycoccus* strains. The number of strains sampled at each site is indicated. Colors correspond to the oceanic basin of origin: blue, Arctic Ocean; green, Atlantic Ocean; red, Mediterranean Sea; grey, Indian Ocean. (B) Summary of the sequencing, assembly and annotation strategy.

*De novo* assembly of the 28 *Bathycoccus sp.* strains resulted in assembly statistics that were either superior to or on par with the reference genome **(Data S1)**. These assemblies have a BUSCO completion score between 94.10% and 97.10%, with a median value of 96.80% and showed an average QPHRED score of 44.28 ± 4.96 (**Figure 2A, 2B)**. These high quality assemblies also displayed an average of 24 contigs per assembly, with a mean N50 of 920 kb, including six assemblies without any identified gap (21 contigs) highlighting their high continuity (**Figure 2C).**

**Fig 2:**
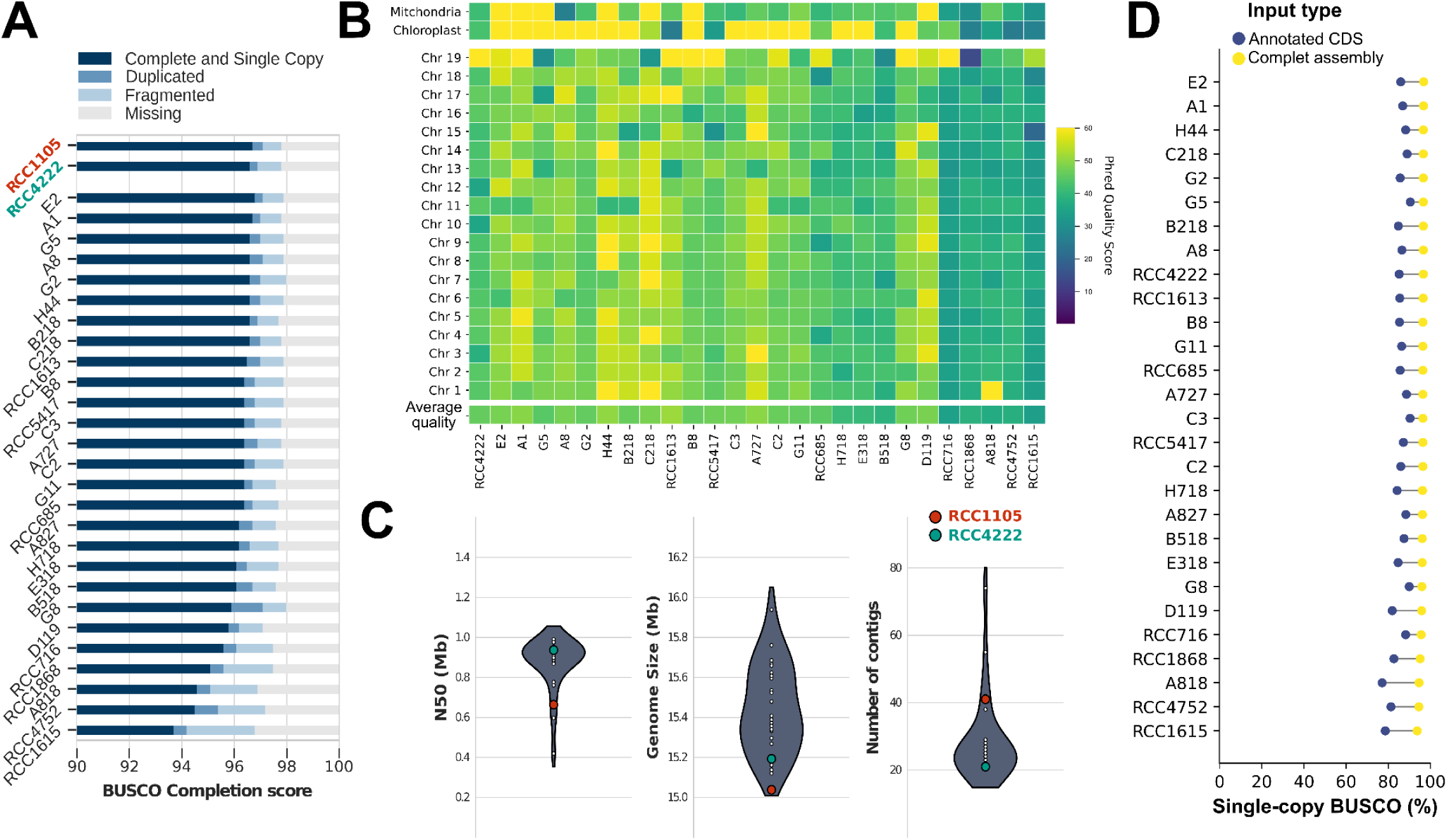
Genome assemblies statistics and quality assessment. (A) BUSCO completion score of genome assemblies for the 28 *de novo* assembled *Bathycoccus* genomes using the *chlorophyta_odb10* library. RCC1105 corresponds to the reference genome produced by Moreau et al. (2012), while RCC4222 corresponds to the *de novo* assembly obtained from a clonal strain of RCC1105. (B) PHRED quality score of genome assemblies per chromosome. (C) Distribution of principal assembly statistics. The N50 corresponds to the sequence length of the shortest contig at 50% of the total assembly length. (D) Comparison of BUSCO single-copy completion score using complete genome sequence or annotated CDS sequences as input.

RCC4222, a clonal strain of the *B. prasinos* reference strain RCC1105 isolated in 2006 (Moreau et al., 2012), was sequenced in 2018 to check clonal conservation of the genome structures and to test the assembly pipeline. No major structural variations were identified compared to the original reference **(Figure S2)**. The chromosome 19, a small outlier chromosome (SOC) described in *Bathycoccaceae* as an hypervariable structure with a high intraspecific diversity, was fully conserved between RCC4222 and RCC1105 (Moreau et al., 2012; Blanc-Mathieu et al., 2017). This indicates that the *B. prasinos* genome is stable in culture and validates our pipeline. Overall, reassembly of the reference using long reads improved continuity, as evidenced by the higher N50 (from 663 to 937 kb), higher genome size (from 15.04 to 15.19 Mb), and a reduction of contig number from 41 to 21, corresponding to the 19 nuclear chromosomes in addition to the the chloroplastic and mitochondrial genomes. (**Figure 2C**). The BUSCO completion scores remained comparable between assemblies, confirming furthermore the higher completion of the resequenced reference genome (**Figure 2A**).

A non-redundant repeat library, generated from all *Bathycoccus sp.* genomes with RepeatModeler and CD-hit, was used for repeat annotation *via* RepeatMasker, resulting in the annotation of ∼10.36 ± 0.82% of each genome as repeated sequences **(Figure S3, Table S2)**. However, only a few of them could be associated with known structures, including LINEs, LTR elements and transposons, while the majority of classified sequences corresponded to simple repeats (∼3.22 ± 0.18% of sequences) **(Table S2)**. A structural annotation of coding sequences was performed for each genome using available RNAseq data from *B. prasinos* strain RCC4752 and plant orthologous protein data to train three gene prediction models through the BRAKER3 and Maker pipelines (GENEMARK-ETP, Augustus and SNAP). The output of each model was then integrated using the Maker annotation pipeline into a complete structural annotation (**Figure 1B, Figure S2)**. The structural annotation predicted 7,431 (± 236) genes, with 7,243 annotated in the strain RCC4222 while 7,847 coding genes were initially annotated in the reference (Moreau et al., 2012). This discrepancy may be due to a lower ratio of mono-exonic genes (73% of predicted transcripts) in our annotation, resulting from the integration of genome assembled transcripts from RNAseq data **(Table S3)**. For quality control of the produced annotation, single copy BUSCO completion scores were performed either for complete assembly genome sequences or for protein sequences extracted from annotated CDS (coding DNA sequences). Complete assembly sequences produced an average BUSCO completion score of 96.10 ± 0.75% while annotated CDS produced an average score of 85.72 ± 3.24% (**Figure 2D**), confirming the quality of the structural annotation.

### Phylogenomic of the *Bathycoccus* genus

Using this high-quality dataset, we investigated the phylogenomics of the *Bathycoccus* genus. Maximum likelihood phylogenetic distances were inferred from nucleic acid alignments of genes shared between all strains. To avoid potential annotation biases between species caused by the predominance of *B. prasinos* transcriptomic data in the annotation model training, a set of 1,200 shared BUSCO genes coming from the *chlorophyta_odb10* database were used. The inferred phylogenetic tree showed a clear separations between strains of *B. prasinos*, *B. calidus* and the three putative *Bathycoccus* B3 genome sequences from the Banyuls bay. As expected these putative *Bathycoccus* B3 grouped with the B3 strain UST710 isolated in Hong Kong by Xu et al. (2024), showing an early divergence in the phylogenetic tree (**Figure 3A**). Cell ultrastructure determined by electronic microscopy revealed characteristic features of the *Bathycoccus* genus with a single mitochondria, a single chloroplast containing a single starch granule, and scales at the cell surface that have been described earlier for *B. prasinos* (Moreau et al., 2012), *B. calidus* (Bachy et al., 2021) and *Bathycoccus* B3 strains from Hong Kong (Xu et al., 2024) (**Figures 3B, 3C, 3D)**.

**Fig 3:**
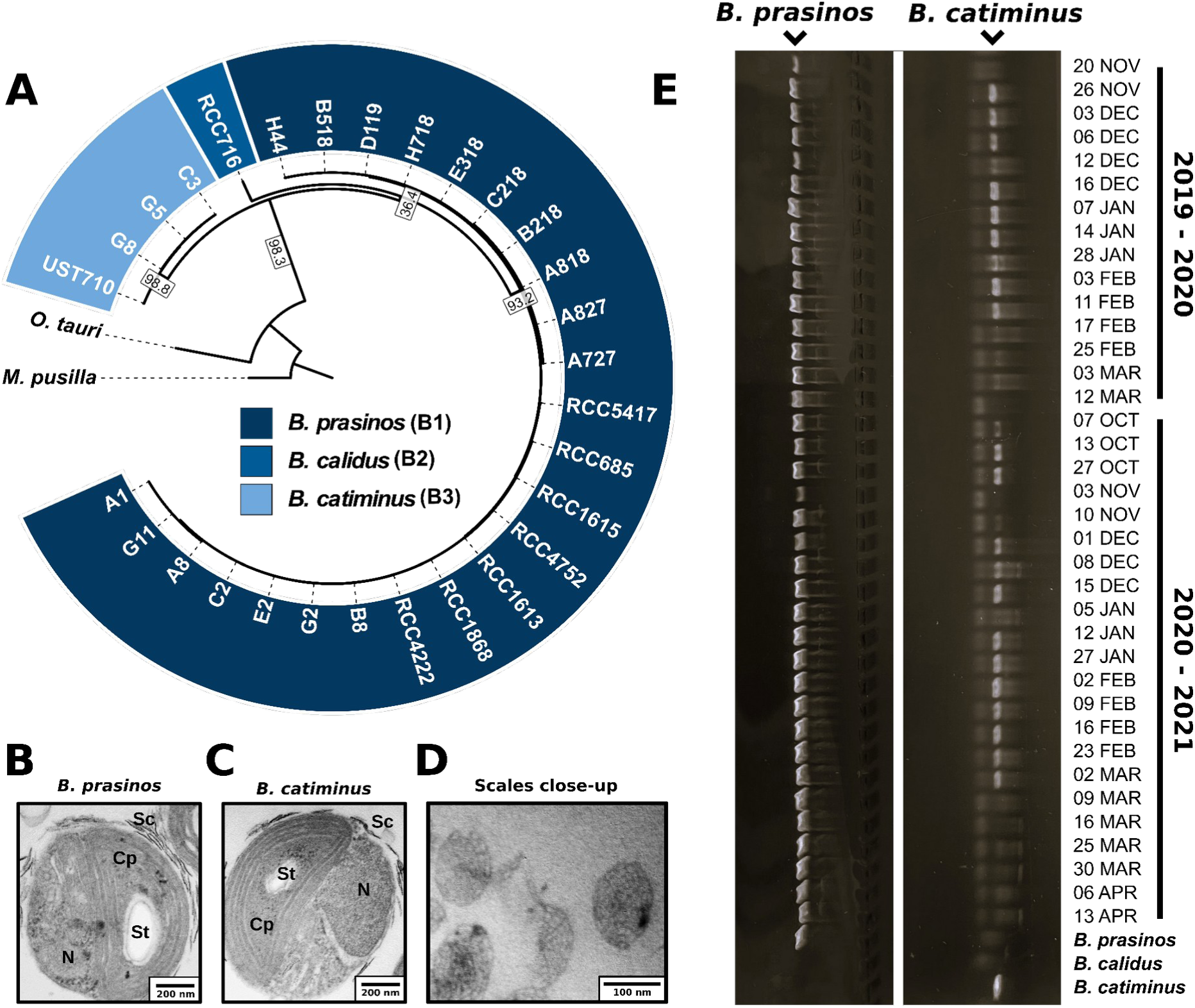
Phylogenomic of the *Bathycoccus* genus and *B. catiminus* characterization. (A) Maximum-likelihood phylogenetic tree of *Bathycoccus* strains based on 1,200 shared genes. Species separation is derived from the branch length and marked by colors. The tree includes strain UST710 isolated and sequenced by Xu et al. (2024) for reference. *O. tauri* and *M. pusilla* are used as outgroups. Only branches with bootstrap values higher than 90% are displayed. Gene concordance factors are displayed for all visible branches. Transmission electron microscopy images of (B) *B. prasinos*, (C) *B. catiminus* strain isolated from the Banyuls bay, and (D) a close up of detached spider web scales from *B. catiminus*. N: nucleus, Cp: chloroplast, St: starch granule, Sc: scales. Bar = 100 nm. (E) Presence of *B. catiminus* in Banyuls bay seawater samples from 2019-2020 and 2020-2021 phytoplanktons blooms, identified using specific primers designed from non-conserved sequences of TOC1 ORF specific to *B. catiminus*.

The mapping of metagenomic data from TARA Ocean and TARA Polar Circle cruises were performed on a representative genome for each species. This analysis confirms the clear latitudinal and depth separations between *B. prasinos* and *B. calidus* populations, initially described by Vannier et al. (2016), with *B. prasinos* found in temperate regions at the surface, and *B. calidus* in tropical water at the deep chlorophyll maximum (DCM). *B. prasinos* was also abundant (up to 4% of sequences) in Arctic waters. *Bathycoccus* B3 was also detected in metagenomic data, and shows geographical distribution patterns similar to *B. prasinos*, however at much lower levels of abundance than *B. prasinos* and *B. calidus* (up to 0.5% of sequences). However, the comparison of mapped metagenomic reads indicates a high level of cross-mapping between *B. prasinos* and *Bathycoccus* B3, with only ∼9% of reads being specific to *Bathycoccus* B3 in all stations. This results in approximately 0.5% of horizontal coverage distributed along all chromosomes of *Bathycoccus* B3 and suggests that its abundance is ∼10 times lower than initially computed **(Figure S4)**. Based on the current evidence of the phylogenetic and spatial-temporal distinctions of the Bathycoccus B3 clade strains, we propose to name this cryptic species as *Bathycoccus catiminus* (from the French expression “en catimini’’ meaning “on the sly”) in reference to its unexpected discovery and its very low abundance in metagenomic datasets compared to *B. prasinos and B. calidus* species **(Appendix S1)**.

The presence of *B. catiminus* was also experimentally investigated through PCR in water samples from the Banyuls bay between January and March 2019, using species specific primers designed from sequences of the *B. catiminu*s TOC1 ORF. *B. catiminus* was detected in Banyuls bay water between November 2019 and February 2020, and between October 2020 and March 2021 (**Figure 3E**), in addition to its presence in December 2018 when the *B. catiminus* strains were isolated.

### Comparative quality assessment of *Bathycoccus prasinos* MAGs and *de novo* genome assemblies from isolated strains

The *B. prasinos* genomic resource was expanded by adding 11 metagenome-assembled genomes (MAGs) available in the literature, as reported by Xu *et al*. (2024), to our 24 *de novo* assembled genomes to investigate the species’ intraspecific phylogenetic diversity. Based on 265 BUSCO genes conserved between the 35 samples, this analysis revealed an apparent disconnection between these datasets, as illustrated by extreme phylogenetic distances among MAGs, and between MAGs and *de novo* assembled genomes (**Figure 4A**). MAGs appeared to cluster together rather than with strains isolated from the same geographical basins, with the exception of TARA_ARC_108_MAG_00264 which presented a weak clustering with other Arctic strains. Moreover, comparison of assembly metrics highlighted drastically lower values for MAGS with a wide distribution compared to *de novo* assembled genomes. For instance, the median N50 values were 0.01 Mb for MAGs compared to 0.92 Mb for assembled genomes (**Figure 4B**). Similarly, the median genome size was approximately 15% smaller in MAGs (13.07 Mb) compared to *de novo* assembled genomes (15.39 Mb) (**Figure 4C),** while the median number of contigs reached up to 1,998 in MAGs for only 23 in *de novo* assembled genomes (**Figure 4D**). Finally, the median BUSCO completion score in MAGs was 85.5% compared to 96.4% in *de novo* assembled genomes (**Figure 4E**). Collectively, these differences highlight the large variations in continuity, contiguity, and completeness among MAGs and compared to *de novo* assembled genomes, limiting their use in high resolution intraspecific diversity characterization including in phylogenetic analysis.

**Fig 4:**
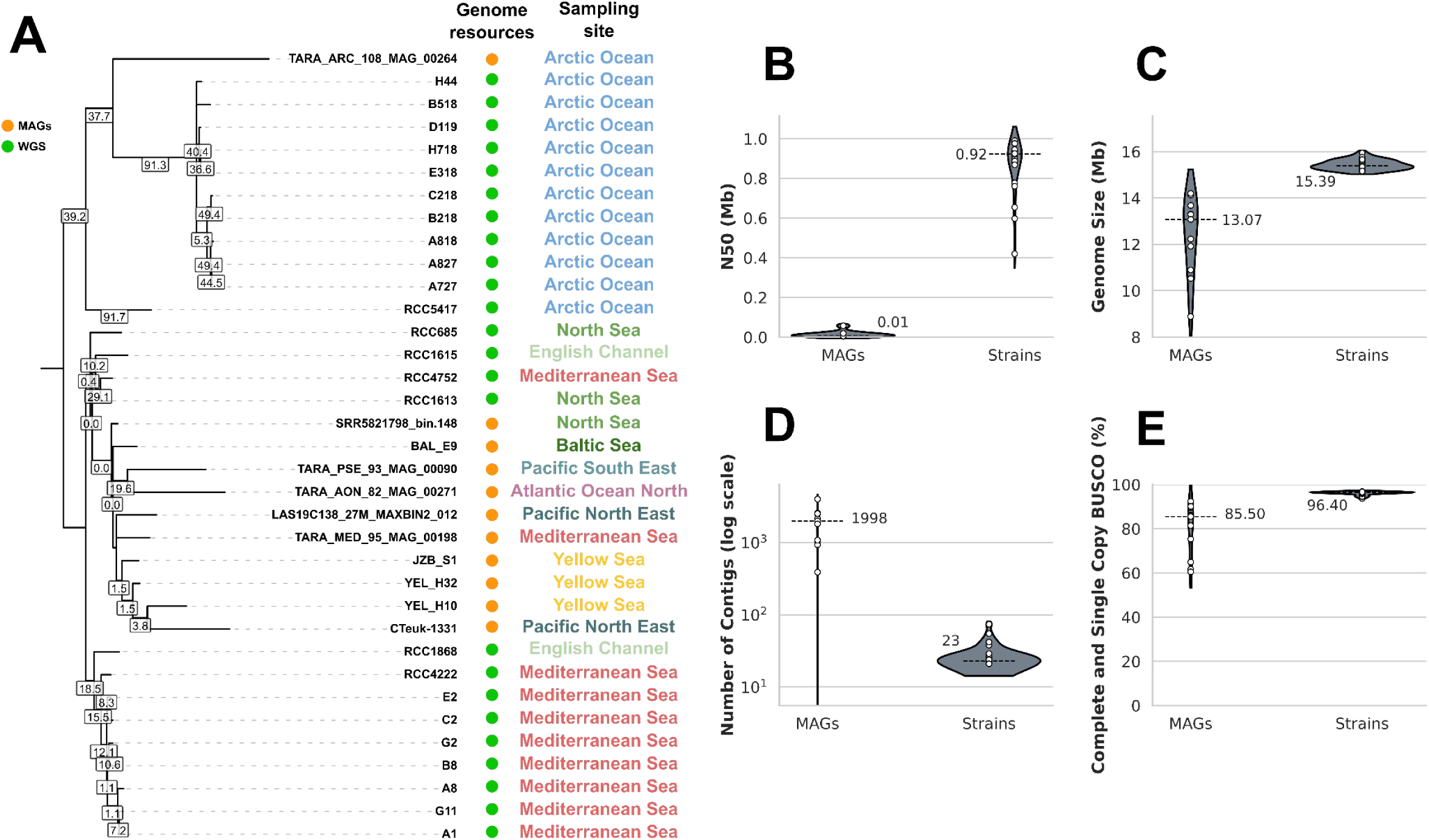
Qualitative comparison between *de novo* genome assembly from whole-genome sequencing data (WGS) and metagenome assembled genomes (MAGs) of *Bathycoccus prasinos*. (A) Maximum-likelihood phylogenetic tree of *B. prasinos* available genomes based on 265 shared genes. Only branches with bootstrap values higher than 90% are displayed. Gene concordance factors are displayed for all visible branches. (B to E) Distribution of principal assembly statistics between available MAGs (n=11) and whole genome assemblies produced from isolated strains of *B. prasinos* (n=24). Median values are displayed for reference.

### Intraspecific diversity and chromosomal haplotypes in *Bathycoccus* strain genomes

For an accurate assessment of *B. prasinos* intraspecific diversity, the phylogenetic diversity of the *Bathycoccus prasinos* species was inferred focusing on the 24 corresponding sequenced genomes and using 1,200 conserved genes. The resulting phylogenetic tree revealed two main branches, separating the strain isolated from the North of Baffin bay during the DarkEdge cruise in October 2021 from all other strains, including the arctic strain RCC5417 isolated from the south of Baffin bay in 2016. However, this last strain diverges early from the cluster of temperate strains. Similarly, Mediterranean strains from the Banyuls bay isolated in 2018 and 2019 were more closely related to each other than to the RCC4222 strain isolated from the bay of Banyuls in 2006. Comparably, the strain RCC4752 isolated in 1986 in the Gulf of Naples did not cluster together with other Mediterranean strains. Geographical origin of strains from the English Channel and the North sea were not resolved in our phylogenetic tree (**Figure 5**). Within geographic basins, subgroups were observed including 3 strains in the Banyuls bay (A1, G11 and A8) and 5 strains in the Baffin bay (C218, B218, A818, A827, A727). A second phylogenetic analysis was performed based on 69 shared genes located on chromosome 14, the BOC of *B. prasinos*. This analysis revealed that strains within the aforementioned subgroups clustered together, independently of their geographic origin. Furthermore, using the same approach, the sister species *B. calidus* and *B. catiminus* clustered together with *B. prasinos* regardless of their taxonomic classification, revealing that the two BOC haplotypes are shared within the *Bathycoccus* genus. These were designated as BOC A and BOC B (**Figure 5**).

**Fig 5:**
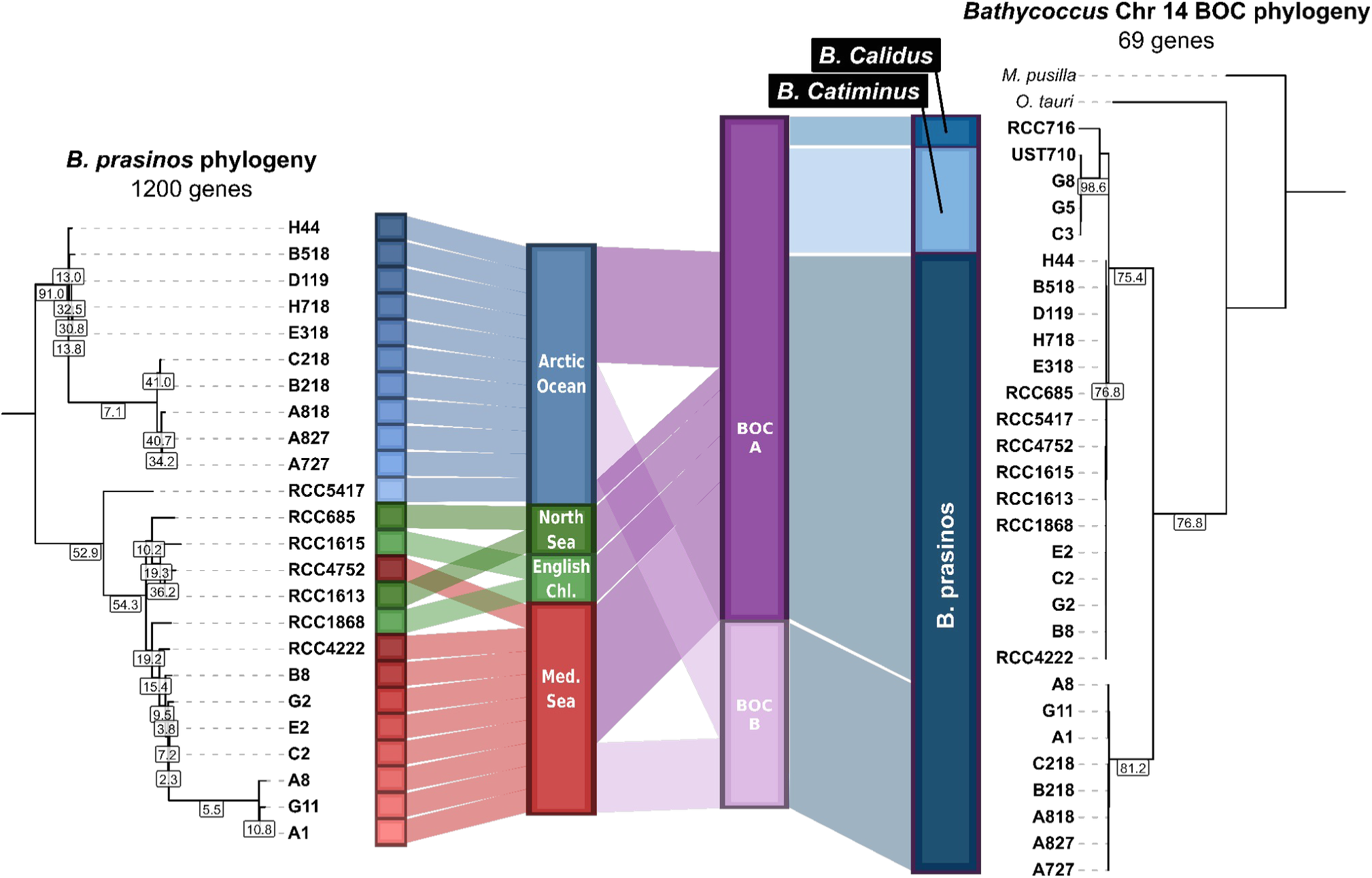
Phylogenomic of *B. prasinos* linked to strain origin and Chromosome 14 BOC haplotypes. (Left) Maximum-likelihood phylogenetic tree of *B. prasinos* strains based on 1,200 shared genes. (Right) Maximum-likelihood phylogenetic tree of *Bathycoccus* strains based on 69 shared genes located on chromosome 14. Only branches with bootstrap values higher than 90% are displayed. Gene concordance factors are displayed for all visible branches.

### Genomic diversity and geographic distribution of big outlier chromosome

The genomic diversity between BOC haplotypes of chromosome 14 was investigated by alignment of chromosome sequences between strains. The identification of syntenic regions confirms that the haplotype specific sequences are restricted to the outlier region, featured by a lower GC content. This chromosome segment corresponds to a ∼400 (BOC A) to 500 kb (BOC B) non-syntenic region between the two haplotypes (*i.e.* ∼2/3 of the chromosome length), with major inversion and duplication events, as well as other smaller structural rearrangements (**Figures 6**). Since several strains were sequenced for each BOC haplotype (16 for BOC A, 8 for BOC B), single nucleotide polymorphism as well as structural variation density along chromosome 14 could be determined. Polymorphism density was computed on a 20 kb sliding window along chromosome 14, and compared to the genome-wide polymorphism to detect local divergences in sequence diversity. Lower than average polymorphism (local density/average density < 1) was detected in the outlier region of both identified haplotypes and, overall, lower polymorphism density of this region could be seen compared to the common regions located at both extremities of chromosome 14 (**Figures 6B, 6C, Figure S5)**. This clear pattern of localized reduced polymorphism could not be seen on any other non outlier chromosomes **(Figure S6)**.

**Fig 6:**
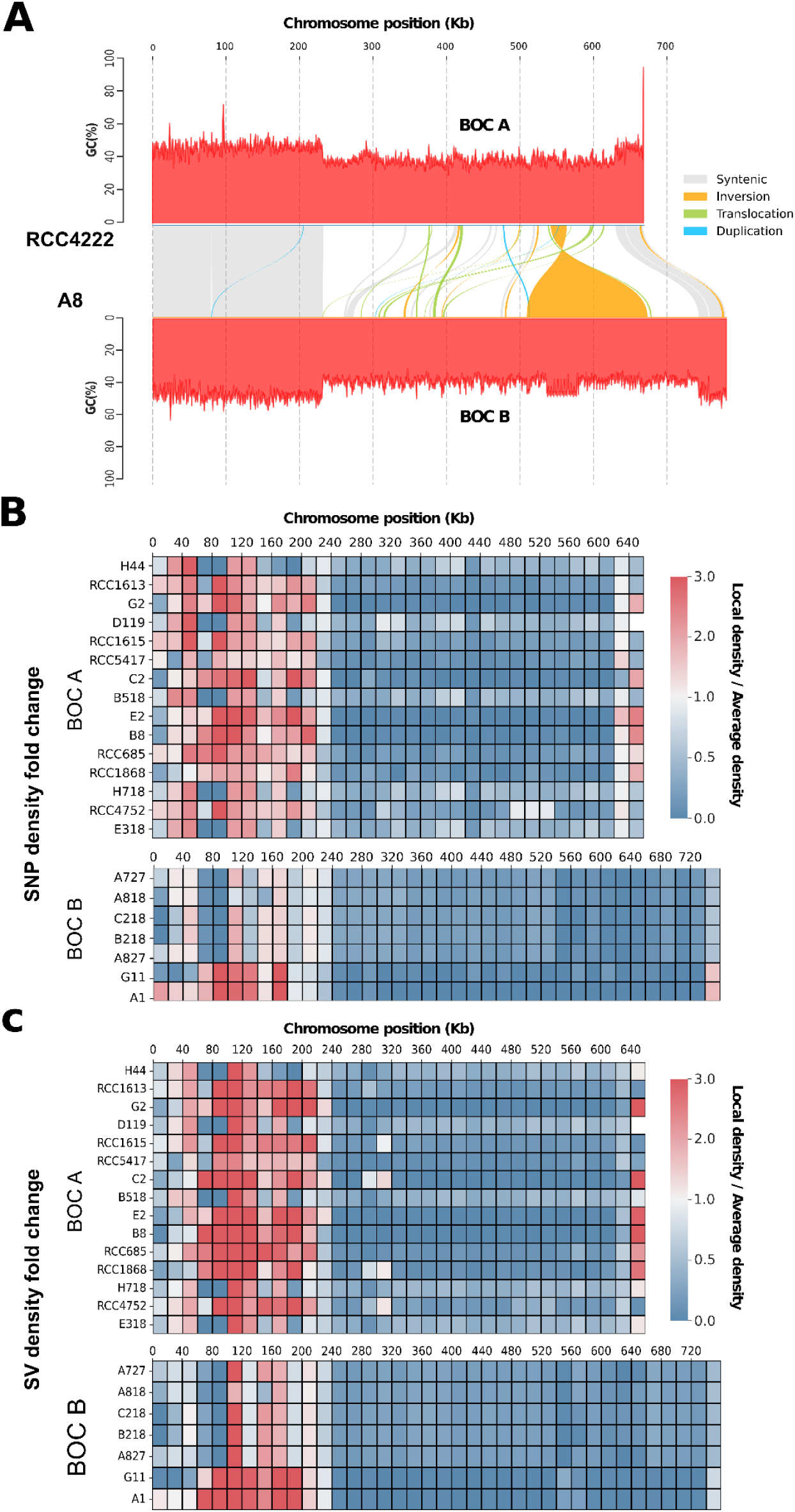
Genomic diversity and haplotypes biogeography of *Bathycoccus prasinos* big outlier chromosome. (A) Alignment of Chromosome 14 from *B. prasinos* strains RCC4222 and A8. Colors indicate syntenic regions and structural rearrangements. Y axis displays the GC content of the corresponding sequences. (B) Divergences between local single nucleotide polymorphism (SNP) density and genome wide SNP density along a 20 kb sliding window. (C) Divergences between local structural variation (SV) density and genome wide SV density along a 20 kb sliding window. SNP and SV calling was computed for each BOC haplotype against strain RCC4222 for BOC A and strain A8 for BOC B.

The geographical distribution of both BOC haplotypes in the world Ocean was assessed by mapping of TARA Ocean and TARA polar circle metagenomic dataset on their respective outlier sequences. In all stations defined by relatively abundant *B. prasinos* sequences, the BOC A haplotype, corresponding to the haplotype of the reference strain RCC4222, was predominant, with a mapping ratio ranging from 68% to 97% of all mapped reads. This imbalance seemed less strict in polar circle stations, with the BOC B haplotype representing up to 32% of mapped reads (**Figure 7A**). This difference in BOC ratios between temperate and arctic regions was further amplified in the haplotype ratio of isolated strains, with BOC A corresponding to the majority in strains isolated from the Banyuls bay (63.6% of strains), while BOC B was predominant in the Baffin bay (82% of strains) (**Figure 7B**).

**Fig 7:**
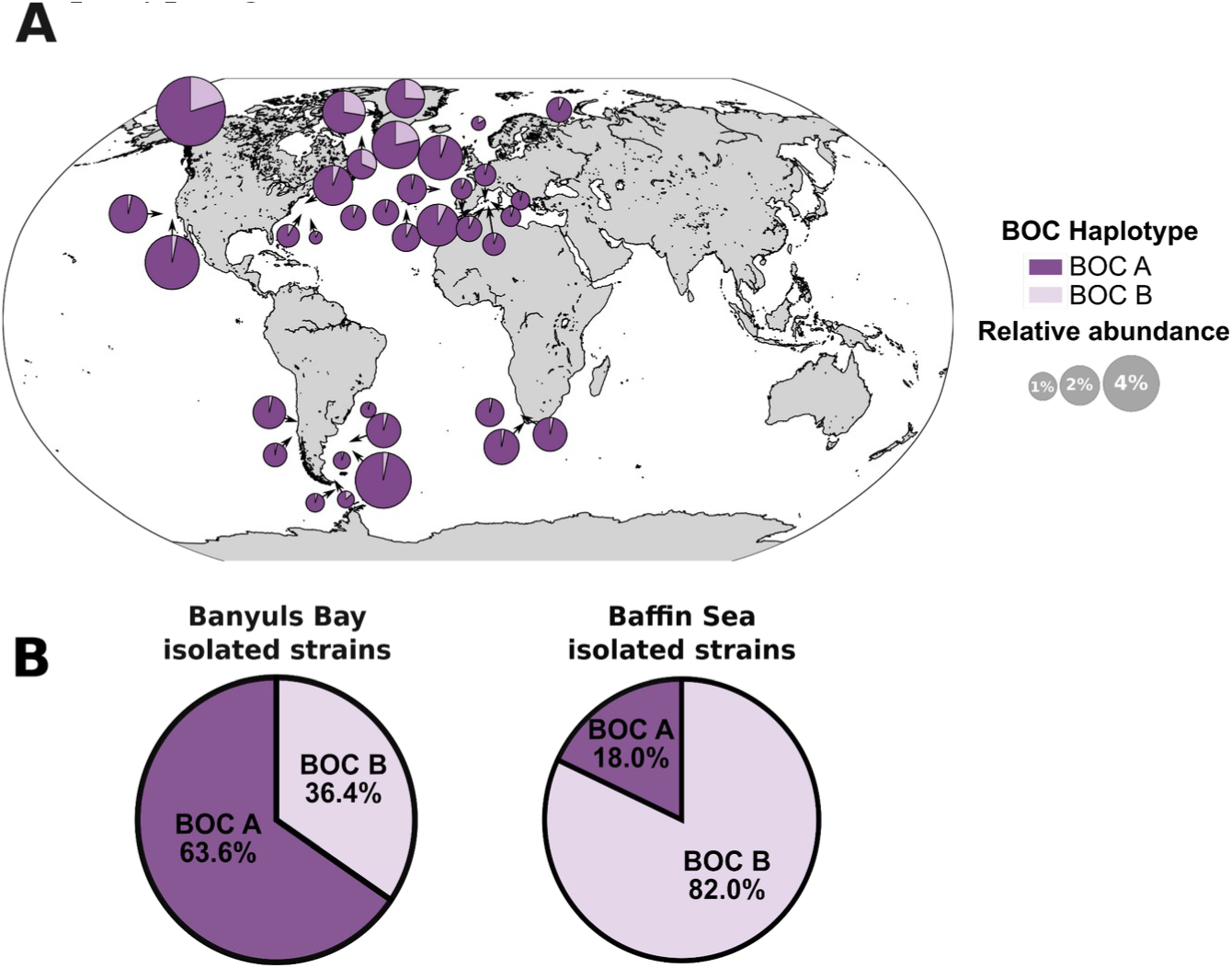
Characterization of BOC haplotype geographical distribution. (A) BOC haplotypes ratio based on read mapping from TARA ocean metagenomic datasets in stations with *B. prasinos* relative abundance > 0.1%. (B) BOC haplotype ratio among strains isolated in the Banyuls Bay (Left)(n=55) and the Baffin Sea (Right)(n=183).

### Genomic diversity of the small outlier chromosome

Another characteristic feature of *Mamiellophyceae* genomes is the small outlier chromosome (SOC), corresponding to chromosome 19 in *Bathycoccus* species. The SOC chromosome size varied considerably among the 24 assemblies of *B. prasinos*, particularly in arctic strains, with sizes ranging from 48 to 230 kb. With the exception of RCC1868 from the English channel, for which major conserved and syntenic regions were identified with strains A8, G2 and A818 (up to 190 kb conserved with A8), little or no synteny was detected between SOCs (**Figure 8, Figure S7)**. This held true even between strains originating from the same geographic basin or isolated at the same time in both Banyuls and Baffin bays. Surprisingly, despite the low synteny, conserved sequences with a wide range of sizes (Average size : 15,787 ± 22,706 bp) were identified, indicating extensive reorganization within each genome (**Figure 8**).

**Fig 8:**
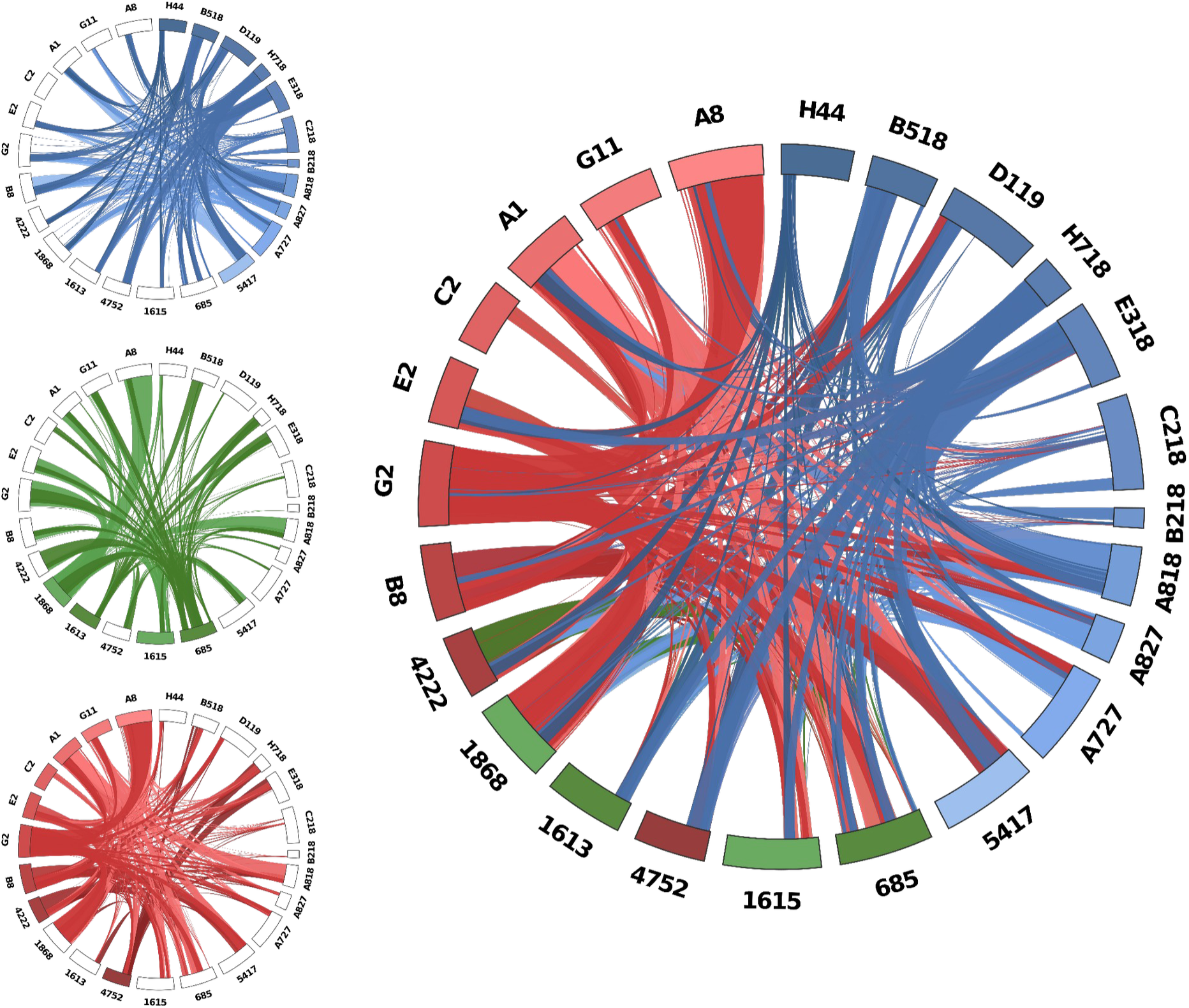
Chromosome 19 SOC shared sequences between *B. prasinos* strains. Chord plot representation of chromosome 19 shared sequences between strains of the geographically close water basins (Left) and between all 24 *B. prasinos* sequenced strains (Right). Chromosome color corresponds to basin and geographical distribution of strains as shown in Figure 5 : Blue, Arctic ocean ; Green, North sea and English channel ; Red, Mediterranean sea. Only shared sequences longer than 500 bp are displayed.

Higher chromosome 19 sequence conservation was observed between strains isolated from the same geographic region, at the same time, such as for the Baffin bay in 2021, or the Banyuls bay in 2018-2019, up to the point that several assemblies had their chromosome almost fully composed of shared sequences (**Figure 8**). However, chromosome 19 of Mediterranean strains isolated earlier (such as RCC4752 isolated in 1986 in the Gulf of Naples, or RCC4222 isolated in 2006 in the Banyuls bay) had fewer conserved sequences with chromosome 19 of other Mediterranean strains than with strains from other geographical regions. Apart from the strain C2, which showed no chromosome 19 conservation with arctic assemblies, all genomes had at least one conserved region with another strain (**Figure 8**). Remarkably, all sequences of the shortest chromosomes 19 (H718, B218 and A827 strains) were detected in other strains. These conserved sequences were usually concentrated in a distal region of chromosome 19, as seen in strains from the Banyuls bay and the Baffin bay **(Figure S7)**.

## 5. Discussion

### Worldwide biological and genomic resources for the study of latitudinal and seasonal adaptation

In this study, we isolated 183 Arctic strains to add to the 66 Mediterranean strains previously isolated from the Banyuls bay (Devic et al., 2024), and to the available strains of the Roscoff Culture Collection, further extending the current *Bathycoccus sp.* biological resource. All arctic and mediterranean strains were genotyped, and the main haplotypes were fully sequenced using ONT and Illumina sequencing, including 10 strains from the Baffin bay and 10 strains from the Banyuls bay. This biological resource, covering a latitudinal gradient from pole to equator, provides an unprecedented tool to study the *Bathycoccus sp.* diversity and the physiological responses underlying adaptation to latitudinal gradients and seasons. Phylogenomic analysis revealed that 3 strains isolated from the Banyuls bay corresponded to the recently described *Bathycoccus* B3 clade that we named *Bathycoccus catiminus* (**Figure 3**). Unlike *B. prasinos* and *B. calidus*, *B. catiminus* is barely detectable in large metagenomic datasets such as TARA Ocean and TARA Polar Circle, being restricted to a few geographic locations such as the Yellow Sea and the Caspian Sea (Xu et al. 2024). Concurrently, sequencing of ITS2 rDNA regions in the Kara sea and MAG reconstruction indicates that a fourth species is yet to be isolated (Belevich et al., 2021; Xu et al., 2024) **(Figure S1)**, highlighting the underestimated taxonomic diversity of the *Bathycoccus* genus. As reviewed by Yung et al. (2022), the description of new species following genome sequencing is frequent in *Mamiellophyceae,* due to the lack of polymorphism in cellular ultrastructure and in the sequence of rRNA 18S V4 and V9 regions, commonly used as distinguishing characters (Piganeau et al., 2011).

The exploitation of natural diversity has greatly contributed to the characterization of genomic features and to the exploration of adaptive mechanisms, not only in *Arabidopsis* but also in other wild land plants and crops cultivars (Alonso-Blanco et al., 2016; Gabur et al., 2019 for review). However, the intraspecific diversity of eukaryotic phytoplankton remains widely unexplored, with only a few studies conducted such as in *Ostreococcus tauri* (Blanc-Mathieu et al., 2017), *Phaeodactylum tricornutum* (Rastogi et al., 2020) and *Emiliania huxleyi* (Read et al., 2013; Bendif et al., 2023). In all these studies, reference-assisted assembly approaches based on short reads mapping to a single reference genome were used. This allows for high resolution of single nucleotide polymorphism and small structural variations, but ignores larger structural variations and population-specific sequences. However, larger modifications also play an important role in environmental adaptation, since they may contain additional genes or regulatory sequences (Gordon et al., 2017). The application of *de novo* assembly methods based on long-read sequencing in *Bathycoccus sp.* enables the identification of such large structural variations and the resolution of complex genomic structures specific to this taxonomic class (**Figures 6, 7 and 8).** While the current state of algal genomic data is characterized by assembled genomes of heterogeneous quality and completeness, restricting intraspecific comparisons to a few generally incomplete genomes (Hanschen and Starkenburg, 2020), we aimed to produce a resource for the study of intraspecific genetic diversity within the *Bathycoccus* genus. Thus, we report here 24 *de novo* assembled and annotated genomes of the cosmopolitan species *B. prasinos*. To our knowledge, this is the largest genomic resource associated with a biological resource available for a single species of green algae (Hanschen and Starkenburg, 2020; Shi et al., 2021). Thanks to the abundance of *B. prasinos* in large metagenomic datasets, numerous MAGs have been reconstructed, opening the way to the intraspecific comparison of populations adapted to specific niches (Xu et al., 2024). However phylogenomic comparisons revealed inconsistencies between available MAGs and our *de novo* assembled genomes (**Figure 4**). MAG reconstruction is not free from biases, since MAGs are assembled from binned reads often originating from geographic basins, rather than from single samples, to compensate for low sequencing depth and horizontal coverage. Therefore, the quality of the assembly is strongly influenced by the sequencing depth and the genetic/genomic heterogeneity of the sequenced population, leading to chimeric, fragmented and most of the time incomplete assemblies with frequent contamination sequences embedded within (Meziti et al., 2021). For instance, the BOC B haplotype of *B. prasinos* was not identified in MAGs, probably due to BOC B representing only 3% to 30% of sequences in metagenomic samples (**Figure 7**). For the same reasons, MAGs are probably not well suited to identifying the fine genetic variations associated with environmental parameters, involved in adaptation to specific environmental niches. The biological and genomic resources presented here, and that will be expanded in the future, should make it possible to identify the genetic signatures underlying the adaptation and ecological success of *B. prasinos* through population genetics and genome-wide association studies similar to those conducted in land plants (Alonso-Blanco et al., 2026; Beaulieu et al., 2025).

### Low polymorphism within putative sexual chromosome haplotypes indicates strong selective pressure

We used the *B. prasinos* resource consisting of 24 genomes together with the existing large metagenomics dataset to conduct an analysis of outlier chromosomes. The big outlier chromosome (BOC), discovered in *O. tauri*, is a putative sexual chromosome that has been identified in all other *Mamiellophyceae* genomes (Derelle et al., 2006; Grimsley et al., 2015). While two haplotypes of this BOC were identified in *O. tauri*, only one haplotype was previously described for the *Bathycoccus* genus (Moreau et al., 2012; Blanc-Mathieu et al., 2017). In this study, the 2 BOC haplotypes were identified in *B. prasinos* strains from both the Banyuls and the Baffin bays, suggesting that these haplotypes are not the result of populations segregation due to geographic isolation, but rather a defining feature of *B. prasinos* genome, as it was previously reported for other *Mamiellophyceae* (Grimsley et al., 2015).

The BOC of *B. prasinos*, corresponding to chromosome 14, has been characterized by several distinct features located in an outlier region, including lower GC content, high number of introns, and increased expression levels (Derelle et al., 2006; Moreau et al., 2012). In *O. tauri*, previous reports indicated a higher linkage disequilibrium and reduced recombination events in the outlier region, leading to an accumulation of transposable elements and suggesting a putative mating function of this region (Blanc-Mathieu et al., 2017), thus following the model of the mating chromosome of *Chlamydomonas reinhardtii* (Ferris and Goodenough, 1994). However, no accumulation of transposable elements was detected in the *B. prasinos* reference genome, and no loss of synteny was identified in strains of the same BOC haplotype **(Figure S5)** (Moreau et al. 2012). Instead, a lower density of single nucleotide polymorphism and structural variations was observed in the outlier region of both *B. prasinos* BOC haplotypes, even between geographically distant strains (**Figure 5B**, **Figure 6C**). Lower than average polymorphism has also been reported in *O. tauri* outlier region and *C. reinhardtii* mating chromosome (Blanc-Mathieu et al., 2017; Hasan et al., 2019), in apparent contradiction with recombination suppression that would intensify mutation accumulation in sexual chromosomes. Gene conversion between mating types was proposed as a sequence homogenizing mechanism by De Hoff et al. (2013), but this hypothesis proved difficult to test with *O. tauri* genomic data, due to the representation of one of the putative mating types by just a single strain. In *B. prasinos*, our analysis of shared genes between BOC haplotypes revealed a phylogenetic separation, and suggests a divergent evolution of chromosome 14 shared genes between haplotypes (**Figure 5**). Strong mating-type-specific selective pressure linked to background selection, as suggested by Hasan et al. (2019, 2020), might be responsible for the reduced diversity and strong differentiation observed in this complex genomic context. Furthermore, this pressure may contribute to the preservation of increased gene expression and fragmented gene structure in the outlier region of the BOC. Although the fitness impact of this 400 to 500 kb haplotype-specific region still needs to be assessed, it is possible that the regional imbalances observed in metagenomic dataset in *B. prasinos* (**Figure 5C**) and *Ostreococcus lucimarinus* (Leconte et al., 2020) could be attributed to either differential fitness, or seasonal dynamics among BOC haplotypes, or both. The cosmopolitan distribution of *B. prasinos*, along with its abundance in metagenomic dataset and the important biological resource available, offers a unique opportunity to monitor the spatio-temporal population dynamics of BOC haplotypes, and to further study the role of BOCs in an environmental context (Vannier et al., 2016; Lambert et al., 2019).

### The viral immunity chromosome presents hypervariable size and a bipartite chromosome structure in the environment

The small outlier chromosome (SOC) is a hypervariable chromosomal structure common to *Mamiellophyceae* genomes, and characterized in *O. tauri* by a low synteny, a high sequence diversity between strains due to extensive structural rearrangements and an important variation in chromosome size (Derelle et al., 2006, Blanc-Mathieu et al., 2017). In this study, we report SOC conserved and rearranged sequences between 24 *B. prasinos* strains. This genomic resource encompasses broad geographical and temporal scales, revealing significant sequence conservation between strains from separate oceanic basins, despite the overall lack of synteny. While several strains come from the same water samples in the Baffin and Mediterranean sea, others were isolated at different times over a period spanning from 1986 to 2020, offering the opportunity to study both the diversity and dynamics of SOC structure at different spatial and temporal scales.

Important size variations for the SOC were displayed in our dataset, especially between Arctic strains isolated during the DarkEdge cruise for which we identified the smallest SOCs (**Figure 6**). Structural and transcriptional modifications of the SOC were shown by Yau et al. (2016, 2018) to be induced by prasinovirus infection and linked to increased spectrum of resistance, suggesting a role of the SOC in viral resistance mechanisms in *Mamiellophyceae*. Moreover, switches between resistant and susceptible phenotypes within isogenic cultures were associated with SOC rearrangements in *O. tauri* and *O. mediterraneus,* and proposed as a coexistence mechanism in microalgae-virus interactions (Yau et al., 2020). As shown by Blanc-Mathieu et al. (2017), smaller SOC sizes are associated with decreased resistance spectrum. Thus, the observed size variations, particularly in arctic strains, might reflect the immune history of the corresponding strains, selecting a more compact chromosome 19 for better local fitness and specific resistance at the expense of broader resistance range. However, these could also be the sign of continuous rearrangement at the time of sampling due to ongoing viral infection.

The smallest SOCs may also provide valuable insight into the structure of *B. prasinos* chromosome 19, as the sequences found in these reduced chromosomes tend to be conserved in nearly all strains and located at a distal position. This separates the SOC into a more conserved portion and a more variable one, with the smallest chromosomes potentially representing a minimal chromosomal content. This observation aligns with the bipartite pattern of SOC transcription reported for *O. tauri* by Yau et al. (2016), in which one half of the chromosome is expressed in susceptible strains while the other half is expressed by resistant strains. Therefore it suggests a common regulation of both transcription and structural rearrangement in this region to minimize loss of fitness in favor of genomic diversity for rapid genomic adaptation. Yau et al. (2016) proposed several theories to explain these observations, including the activation of retrotransposons and epigenetic modifications in response to biotic stress. To comprehensively test these theories, an in-depth characterization of SOC structural variations, including gene-content and structure, is necessary. As such, the extensive dataset described here represents a valuable resource for understanding the intricate interactions between *Mamiellophyceae* and associated viruses.

### Conclusion

The biological and associated genomic resources of *Bathycoccus* presented in this study cover an unprecedented range of latitudes for a eukaryotic phytoplankton species. Genotyping and *de novo* whole-genome sequencing of newly isolated strains provide high-quality reference genomes for three cryptic species of *Bathycoccus* including, *B. prasinos*, *B. calidus*, and *B. catiminus*. This includes 24 high-quality genomes of the cosmopolitan species *B. prasinos*, allowing to deeply explore the intraspecific diversity of this ecologically significant species.

This resource complements the metagenome-assembled genomes (MAGs) currently available and described by Xu *et al*. (2024). However, comparison of these MAGs with genomic data from isolated strains revealed that while MAGs are crucial for characterizing uncultured species, such as *Bathycoccus* B4 (Xu et al., 2024), their quality is inadequate for resolving genomic structural variations such as for BOC and SOC outlier chromosomes. Moreover, their large variations in completeness and quality limit high-resolution diversity analysis. This emphasizes the importance of isolating and extensively sequencing additional strains of ecologically relevant species, such as *B. prasinos*.

Comparative analysis focused on *B. prasinos* strains also provided further biological insights into the putative function of outlier chromosomes. In addition to identifying a second BOC haplotype that was not detected in MAGs, we observed a low polymorphism in the outlier region of this putative sexual chromosome, suggesting a strong selective pressure consistent with its proposed biological function (Grimsley et al., 2015 ; Blanc-Mathieu et al., 2017). Meanwhile, a more detailed characterization of SOC hypervariability revealed a significantly higher level of sequence conservation than previously reported and uncovered a bipartite SOC structure that varies across oceanic regions. This provides a valuable resource for investigating the molecular mechanisms underlying viral resistance in natural environments.

Altogether, the *Bathycoccus* genomic resources established here pave the way for multi-scale comparative analyses at the genus level, and for the construction of a first species-specific pangenome for *B. prasinos.* This genomic resource, combined with metagenomic studies, physiological analysis of *Bathycoccus* natural variants, and recently developed genetic engineering tools for *B. prasinos* (Faktorova et al., 2020), offer new opportunities to study the molecular basis of adaptation underlying the ecological success of the *Bathycoccus* genus and its cosmopolitan representative species *B. prasinos*.

## 6. Experimental procedures

### Algal strains and culture conditions

Selection of *Bathycoccus sp.* strains from the Roscoff Culture Collection (RCC) and isolated strains from the Banyuls bay are detailed in Devic et al. (2024). Culture conditions of available and isolated Mediterranean strains are described in Devic et al. (2024). Arctic isolates were grown at 4°C or 15°C under constant light (10 µE).

### Sampling and cell isolation

Sea water sampling at the SOLA buoy in Banyuls bay and the isolation of *Bathycoccus* strains were reported earlier by Devic et al. (2024).

Sampling in the Baffin bay was performed during the DarkEdge campaign in October, 2021 (**Figure 1A**). For Baffin bay samples, 50 ml of sea water was filtered through a 1.2 μm pore-size acrodisc (FP 30/1.2 CA-S cat N° 10462260 Whatman GE Healthcare Sciences) and used to inoculate culture flasks with 10 ml of filtrate each. Sea water was supplemented with vitamins, NaH2PO4, NaNO3 and metal traces at the same concentration as in L1 culture medium. For samples isolated from the DE310 ice station, the salinity of the culture medium was halved by adding 10 ml of mQ water. Antibiotics (Streptomycin sulfate 100 µg/ml) were added to half of the samples to limit bacterial growth. Finally, cultures were incubated on board either at 4° C or 15° C for 10-20 days under constant light before shipping through express carriers to Banyuls-sur-Mer (France) at 4° C or 15° C depending on the culture conditions. In the laboratory, the presence of picophytoplankton was checked using a BD accuri C6 flow cytometer. Cultures containing at least 90% of picophytoplankton were used for plating in low melting agarose 0.21% W/v as described by Devic et al. (2024). Colonies appearing after 10-15 days at 15° C, and up to a month at 4° C, were hand picked and further cultured. Individual *Bathycoccus* cells were obtained from 3 sampling zones at 4° C and 15° C with or without antibiotics **(Table S1)**.

### Genotyping

Genotyping, identification or detection of strains were performed by PCR in 25 or 50 µl final volume containing DNA, REDTaq polymerase 2x master mix (VWR), 200µM final of each primer and water. Amplification began by a pre-denaturating step at 95°C for 3 minutes, followed by 40 cycles (95°C for 20 seconds, annealing at 58°C for 30 seconds, extension at 72°C for 1 minute) and a final extension step for 5 minutes. The PCR products were loaded onto an agarose gel to verify their size and, if necessary for Sanger sequencing, to be purified using NucleoSpin Gel and PCR Clean-up kit (Macherey Nagel, Germany).

The identification of *Bathycoccus sp.* strains from the Banyuls bay was performed through specific amplifications of a 614 bp fragment of the LOV-HK gene (Bathy10g02360) and 18S rDNA followed by Sanger sequencing (GENEWIZ), as reported by Devic et al. (2024). 55 isolates were unambiguously identified as *B. prasinos* by Devic et al. (2024), while 11 strains showed a low amplification signal with the LOV-HK primers but were determined to be *Bathycoccus sp.* by 18S rDNA sequencing results. We amplified the ITS2 region with 5’-GTACACACCGCCCGTCGC-3’ and 5’-ATATGCTTAARTTCAGCGGGT-3’ primers for these 11 strains. Sanger sequencing of the PCR products led to the identification of *B. catiminus*. Primers in the variable region of the Flavodoxin-like gene (Bathy03g02080; 5’-GCAAGAGAAGATTGAGGCGGAA-3’ and 5’-CTCTGCTGCCGCTTTTGCCTCA-3’) were designed for *B. catiminus* to select the most variable isolates for whole genome sequencing. Detection of *B. catiminus* in seawater was done by PCR amplification of the same environmental DNA samples reported in by Devic et al. (2024) with specific primers in the TOC1 ORF, TOCNI5B3 (5’-GGGACCCACCACAGGTTGCTGT-3’) and TOCNI3B3 (5’-TACCGCGAGCAGCAACAGTAGT-3’).

Since LOV-HK primers failed to amplify *Bathycoccus* strains from the Baffin Bay, the identification of *Bathycoccus sp.* strains was performed through specific amplification of a portion of TOC1 ORF (Bathy17g01510) with primer TOCNI5 (5’-AGGGGTTTTTGCAGAAACCGCT-3’) and TOCNI3 (5’-TCTCGCATTTGATTTCGAGTCCA-3’). Intra-species diversity of Baffin bay strains was assessed using 2 markers, a fragment of the flavodoxin-like gene (Bathy03g02080) and the C-terminal region of the TIM gene (Bathy14g30100) (Devic et al., 2024), resulting in the identification of 10 multi loci genotypes **(Table S1)**.

### Ultrastructure of *Bathycoccus* species

Cells were prepared according to Chrétiennot-Dinet et al. (1995). Thin sections were stained with uranyl acetate and lead citrate, and observed with a 7500 Hitachi transmission electronic microscope.

### Genome sequencing and assembly

Isolates from the Baffin bay were cultured at 4°C before DNA extraction, other strains were grown at 15°C. DNA extraction and Oxford Nanopore Technology (ONT) genome sequencing were performed on 28 *Bathycoccus sp.* strains as described in Devic et al. (2024). Raw ONT data were basecalled using Guppy 6.1.2 (https://nanoporetech.com) and the SUP model; reads with a PHRED score higher than 7 were retained for assembly. Quality control was performed with NanoPlot 1.19.0 (De Coster et al., 2018). Illumina pair-end sequencing (GENEWIZ & Novogene) was also performed on the same extracted DNA, yielding between 450 Mb and 2 Gb (∼30-130X coverage) of sequences per strain. Paired-end sequencing of 250 bp long was used for strains from the RCC and the Banyuls bay, while 150 bp paired-end sequencing was used for strains isolated during the DarkEdge cruise.

Genome assemblies were produced using FLYE 2.9 (Kolmogorov et al., 2019) (options: -- nano-hq --genome-size 15m). Assembly polishing with ONT reads was performed using MEDAKA 1.5 (https://github.com/nanoporetech/medaka) (options: -m r941_min_high_g360). Assembly correction with Illumina reads was performed using BWA 0.7.17 (Li, 2013) for read mapping, Samtools 1.9 (Danecek et al., 2021) and PILON 1.24 (Walker et al., 2014), with standard options. Scaffolding and direction of contigs on the reference genome (Strain RCC1105) was performed using RagTag 2.1.0 (Alonge et al., 2021). Unmapped contigs, derived from RagTag scaffolding, were taxonomically assigned using Blobtools 1.1.1 (Laetsch and Blaxter, 2017)**(Data S2)**. Most unmapped contigs were labelled as contaminating DNA, mainly including extracted DNA from bacteria associated in culture with *Bathycoccus*. Contigs assigned to the *Chlorophyta* phylum were either small (< 10kb) or presented inconsistent coverage compared to mapped contigs. Therefore all unmapped contigs were considered contaminating DNA and excluded. Gap filling was performed using TGS-GapCloser 1.0.1 (Xu et al., 2020) with Samtools 1.9 and PILON 1.24. One round of Pilon correction was applied to corrected gaps for consistency with the global assembly correction (options: --p_round 1 --r_round 0 --min_nread 3). Abnormal chromosomal fusions due to assembly error were manually checked using D-GENIES 1.5.0 (Cabanettes and Klopp, 2018) and corrected using samtools (samtools faidx) (Danecek et al., 2021).

Assembly statistics were calculated with Assembly_stats 0.1.4 (https://github.com/MikeTrizna/assembly_stats) and QUAST 5.2.0 (Gurevich et al., 2013). Completion was assessed through BUSCO 5.4.4 (Manni et al., 2021) using the *chlorophyta_odb10* library, and quality was estimated with MERQURY 1.1 (Rhie et al., 2020) with the corresponding Illumina reads as input.

Basecalling and assembly scripts used in this paper are available at https://github.com/LouisDennu/Bathycoccus_GenomeAssemblyAndAnnotation.

### Genome Annotation

Genome assemblies were individually submitted to RepeatModeler 2.0.1 (https://github.com/Dfam-consortium/RepeatModeler) to build strain specific repeat libraries, which were subsequently concatenated in a non-redundant repeat library using CD-hit (Fu et al., 2012) (90% identity and coverage). This library was used with RepeatMasker 4.1.2 (https://github.com/rmhubley/RepeatMasker) on all samples to produce soft masked assemblies.

Available Illumina RNAseq whole transcriptome of *Bathycoccus prasinos* strain RCC4752 (SRX554258) (Keeling et al., 2014) was mapped upon the RCC4752 genome assembly using HISAT2 2.1.0 (Kim et al., 2015), and transcripts inferred using STRINGTIE 1.3.4 (Shumate et al., 2022). Plant orthologous proteins database 10 (Kriventseva et al., 2019) was used as protein evidences for annotation

Training of GENEMARK-ETP 4.71 (Brůna et al., 2023) and AUGUSTUS 3.3.3 (Stanke et al., 2008) *ab initio* prediction models were performed through BRAKER3 2.1.6 (Gotoh, 2008; Lomsadze et al., 2014; Buchfink et al., 2015; Gabriel et al., 2023), using transcriptome mapping data and the plant protein database as evidence. Training of SNAP (Korf, 2004) *ab initio* prediction model was performed after a first round of MAKER 2.31.9 (Cantarel et al., 2008; Campbell et al., 2014), using genome assembled transcripts, available *de novo* assembled transcripts (https://www.ncbi.nlm.nih.gov/Traces/wgs?val=HBMR01) and plant protein database as evidence. GENEMARK-ETP, AUGUSTUS and SNAP prediction models were merged through a second round of Maker to produce *de novo* structural annotation.

Annotation scripts used in this paper are available at https://github.com/LouisDennu/Bathycoccus_GenomeAssemblyAndAnnotation.

### Phylogenetic analysis

BUSCO (Manni et al., 2021) analysis with *chlorophyta_odb10* library was computed for each assembled genome and on *O. tauri* and *M. pusilla* reference genomes to be used as outgroups. This resulted in 1,200 predicted genes shared across all genomes. Amino acid sequences were aligned for each gene using MAFFT 7.520 (options: --auto --maxiterate 1000) (Katoh and Standley, 2013) and converted to nucleic acid alignment using PAL2NAL 14 (Suyama et al., 2006) to ensure accurate coding sequence alignment.

Sequence alignments were concatenated in a partitioned supermatrix using catfasta2phyml.pl script (https://github.com/nylander/catfasta2phyml). Maximum likelihood phylogenetic inferences with gene concordance factors were computed through a partitioned analysis for multi-gene alignments using IQ-TREE 2.2.0.3 (Chernomor et al., 2016; Minh et al., 2020), with ModelFinder for automatic model finding (Kalyaanamoorthy et al., 2017) and 1,000 ultrafast bootstrap (Hoang et al., 2018) (options: -B 1000 -m MFP+MERGE).

### rRNA structure prediction

2D structure prediction of the 18S rRNA ITS2 region was computed using RNAfold (Lorenz et al., 2011) and default parameters. Predictions are displayed using centroid structures.

### Comparative genomics

Complete genomes dotplot comparison were computed using D-GENIES 1.5.0 (Cabanettes and Klopp, 2018) with Minimap2 2.24 (Li, 2018) as aligner and “Few repeats” option.

Genome assembly alignments were performed with MUMmer 3.1 (Kurtz et al., 2004) nucmer (options: --maxmatch -c 500 -b 500 -l 200), alignments were then filtered for identity (>90%) and length (>100bp) using delta-filter (options: -m -i 90 -l 100). Synteny, single nucleotide polymorphism and structural variations were identified from the MUMmer output by SYRI 1.6 (Goel et al., 2019) and visualized with Plotsr 0.5.3 (Geol and Schneeberger, 2022).

### Mapping of metagenomic datasets

Metagenomic reads from the TARA Ocean and TARA Polar Circle (De Vargas et al., 2015) campaigns were mapped to genome sequences using BWA mem 0.7.17 (Li, 2013). Samtools 1.13 (Danecek et al., 2021) was used to recover mapped reads (samtools view, options: -F 4) and to remove duplicates to avoid bias due to PCR artifacts. Using bamFilters (https://github.com/institut-de-genomique/bamFilters), mapped reads were filtered out for low-complexity bases (>75%), high-complexity bases (<30%), coverage (<80%) and identity (<95%). Remaining reads were retained for further analysis. Relative abundance was computed for each sample as the number of reads mapped normalized by the number of reads sequenced.

For *Bathycoccus sp.* species biogeography, complete assembled genomes of strains RCC4222, RCC716 and G8 were respectively used as reference for *B. prasinos*, *B. calidus* and *B. catiminus*. For big outlier chromosome haplotypes, Chr14:232230-630064 genome segment of strain RCC4222 and Chr14:231394-743201 genome segment of strain A8 were respectively used as reference for BOC A and BOC B outlier regions, according to loss of synteny.

## 7. Accession numbers

Whole genome sequences, annotation files and basecalled reads of the strains sequenced in this study are available at the European Nucleotide Archive under project accession PRJEB67444. All strains sequenced in this study have been deposited in the Roscoff Culture Collection (RCC11109 to RCC11132).

## Supporting information

SuppData

## 8. Acknowledgements

The authors acknowledge the DarkEdge 2021 cruise to the Canadian Arctic, coordinated by Marcel Babin on the CCGS Amundsen, with Marie-Hélène Forget and the Dark Edge Genomics team, including Benjamin Bailleul, Chris Bowler, Richard Dorrell, Carole Duchene and Connie Lovejoy. They also acknowledge the ISO 9001 certified IRD i-Trop HPC (member of the South Green Platform) at IRD Montpellier for providing HPC resources that have contributed to the research results reported within this paper. URL: https://bioinfo.ird.fr/ - http://www.southgreen.fr. The authors thank the Roscoff Culture Collection for providing access to collected strains and Ian Probert for helping with the taxonomic description of *B catiminus*. They also thank Marie-Line Escande and the platform BioPIC at the Oceanological Observatory of Banyuls-sur-Mer for electron microscopy service. This work, along with the salaries of LD and JR, was funded by the french Agence Nationale de la Recherche (ANR Clima-Clock, ANR-20-CE20-0024 to FYB, AF and OJ).

The authors report no conflicts of interest.

## Author contributions

FYB, FS and MD designed the study. MD, AF and NJ designed and carried out the sampling. MD, JCL, VV and CM isolated, genotyped strains and produced the sequencing data. LD, JR and FS performed bioinformatic analyses. MD, LD, FS and FYB described the novel species *B. catiminus*. LD, FS and FYB wrote the manuscript. All authors reviewed the final version of the manuscript.

