## Supplementary material for "Biological and genomic resources for the cosmopolitan phytoplankton *Bathycoccus:* Insights into genetic diversity and function of outlier chromosomes": SuppData

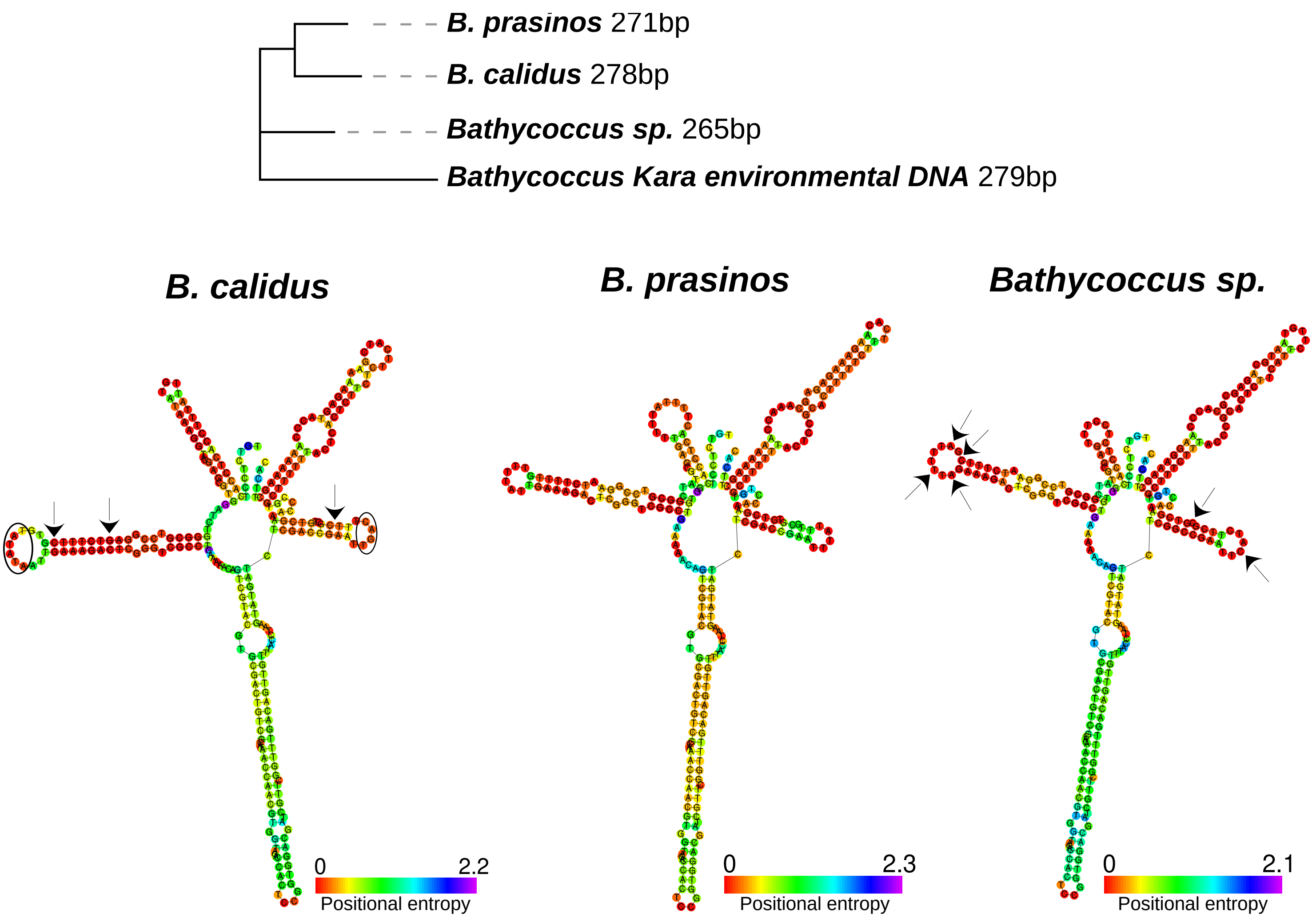


**Fig S1: (A) Maximum-likelihood phylogeny and (B) 2D centroid structure of available *Bathycoccus* 18S rRNA ITS2 region.** Only bootstrap values higher than 90% are displayed. Notable structural variations compared to *B. prasinos* 18S ITS2 region are marked by arrow and circles. 18S sequences were obtained from *B. prasinos* RCC1105 reference strain, *B. calidus* RCC716 strain and *Bathycoccus sp.* G8 strain respectively. *Bathycoccus* Kara environmental sequence was obtained from Belevich et al. (2021).

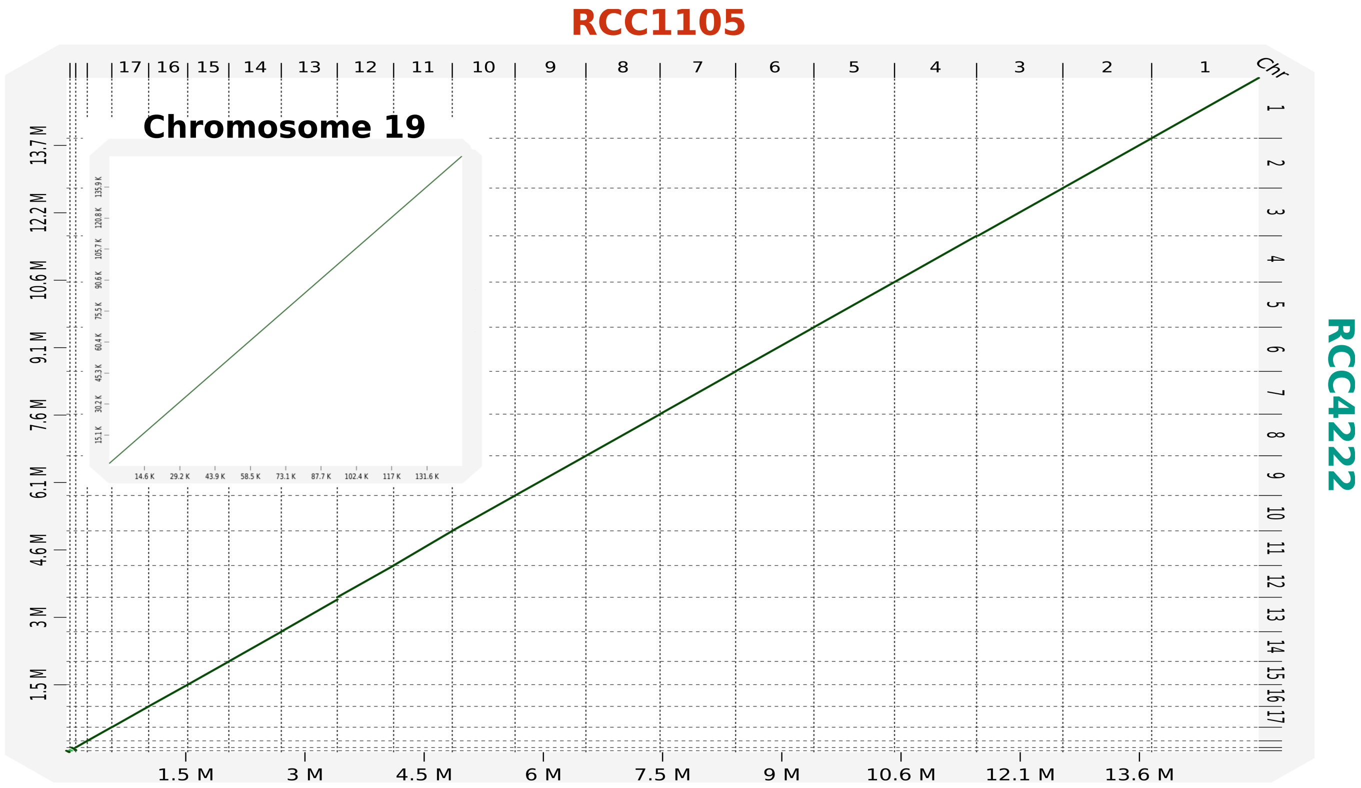


**Fig S2: Clonal conservation of the genome structure between reference strain RCC1105 and clonal strain RCC4222.** Dot-plot of the genome from the reference strain RCC1105 produced by Moreau et al. (2012) against a *de novo* assembled genome of the clonal strain RCC4222 using the assembly pipeline outlined in figure 1B.

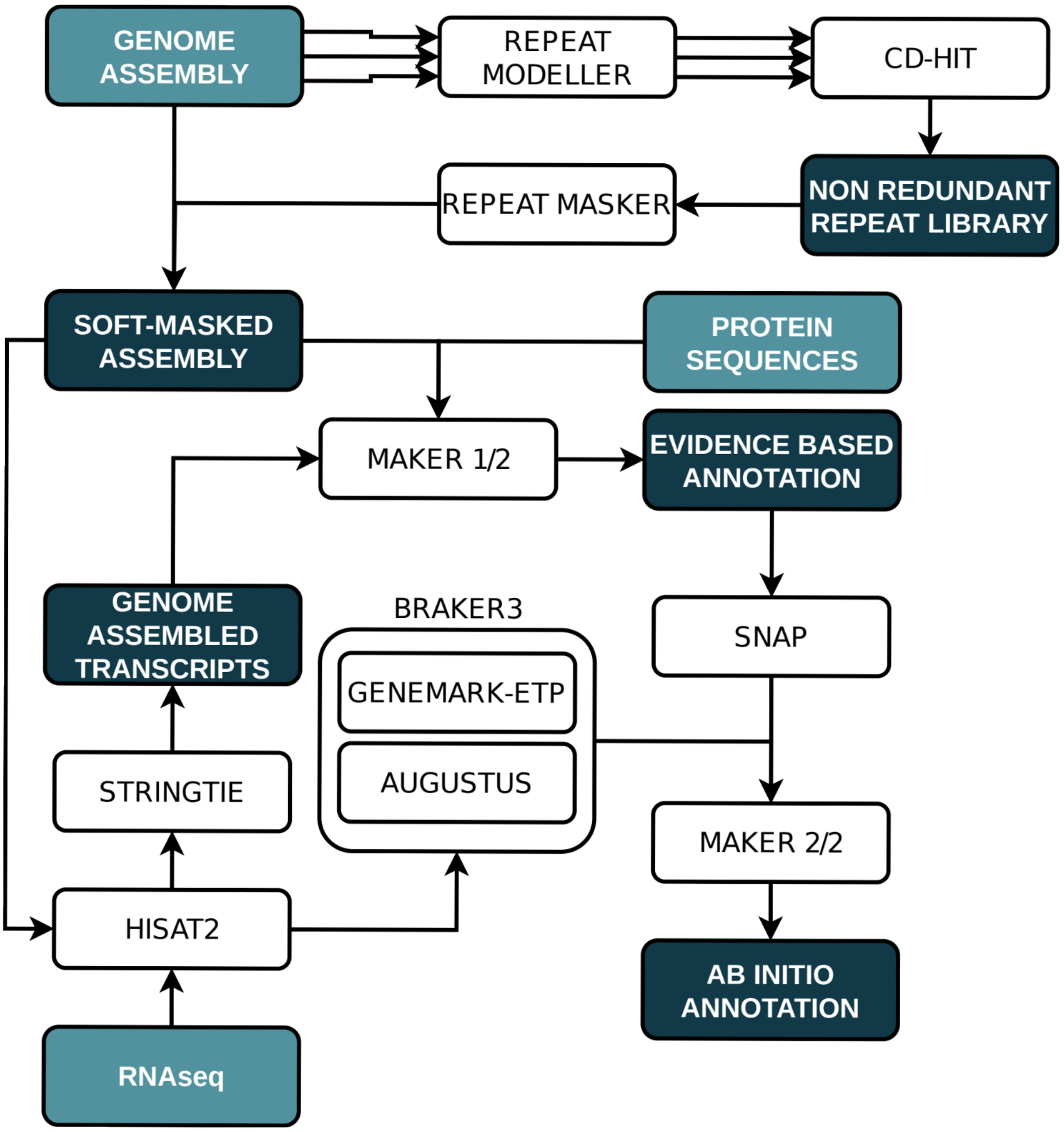


**Fig S3: Structural annotation pipeline of repeat and coding sequences.** Repeat annotation was performed using RepeatMasker and a non redundant repeat library extracted from all 28 *Bathycoccus* genomes assemblies. Gene annotation was performed using RNAseq sequence from strain RCC4752 and protein sequences database as evidences to train Genemark-ETP, Augustus and SNAP models. Models output was integrated using Maker.

**Appendix S1**

**Taxonomic description**

*Bathycoccus catiminus* Dennu, Devic and Bouget sp. Nov.

*Diagnosis*: Morphological characters of the genus (Eikrem & Throndsen 1990). Nucleotide sequences distinct (Whole genome assembly: ENA accession numbers ERZ24811838, ERZ24811841, ERZ24811842). In the second internal transcribed spacer (ITS2) of the nuclear-encoded rRNA transcriptional unit: In helix 2 bases 18 to 20 sequence is 5'-CGA-3'. In the additional helix specific to Bathycoccaceae (Marin et al. 2010) base 12 is C (Figure S1).

*Holotype here designated:* Figure 3B.

*Isotype here designated*: Cryopreserved culture strains here designated as G8, G5 and C3 deposited in the Roscoff Culture Collection (Roscoff Marine Station, France) as RCC11117, RCC11118 and RCC11119 respectively.
*Type locality:* Bay of Banyuls-sur-Mer, Mediterranean Sea (42°28'48"N, -3°31'48"W) at a depth of 3 m.

*Etymology :* from the French word "catimini", meaning hidden, discrete, secret.
*Validating illustration:* Figure 3B.

*Habitat and ecology* : Detected in temperate to high-latitude marine waters and coastal regions at very low abundances.

**Eikrem, W., Throndsen, J.** (1990) The ultrastructure of Bathycoccus gen. nov. and B. prasinos sp. nov., a non-motile picoplanktonic alga (Chlorophyta, Prasinophyceae) from the Mediterranean and Atlantic. Phycologia, 29, 344–350. https://doi.org/10.2216/i0031-8884-29-3-344.1

**Marin, B., & Melkonian, M.** (2010). Molecular Phylogeny and Classification of the Mamiellophyceae class. Nov. (Chlorophyta) based on Sequence Comparisons of the Nuclear- and Plastid-encoded rRNA Operons. Protist, 161(2), 304–336. https://doi.org/10.1016/j.protis.2009.10.002

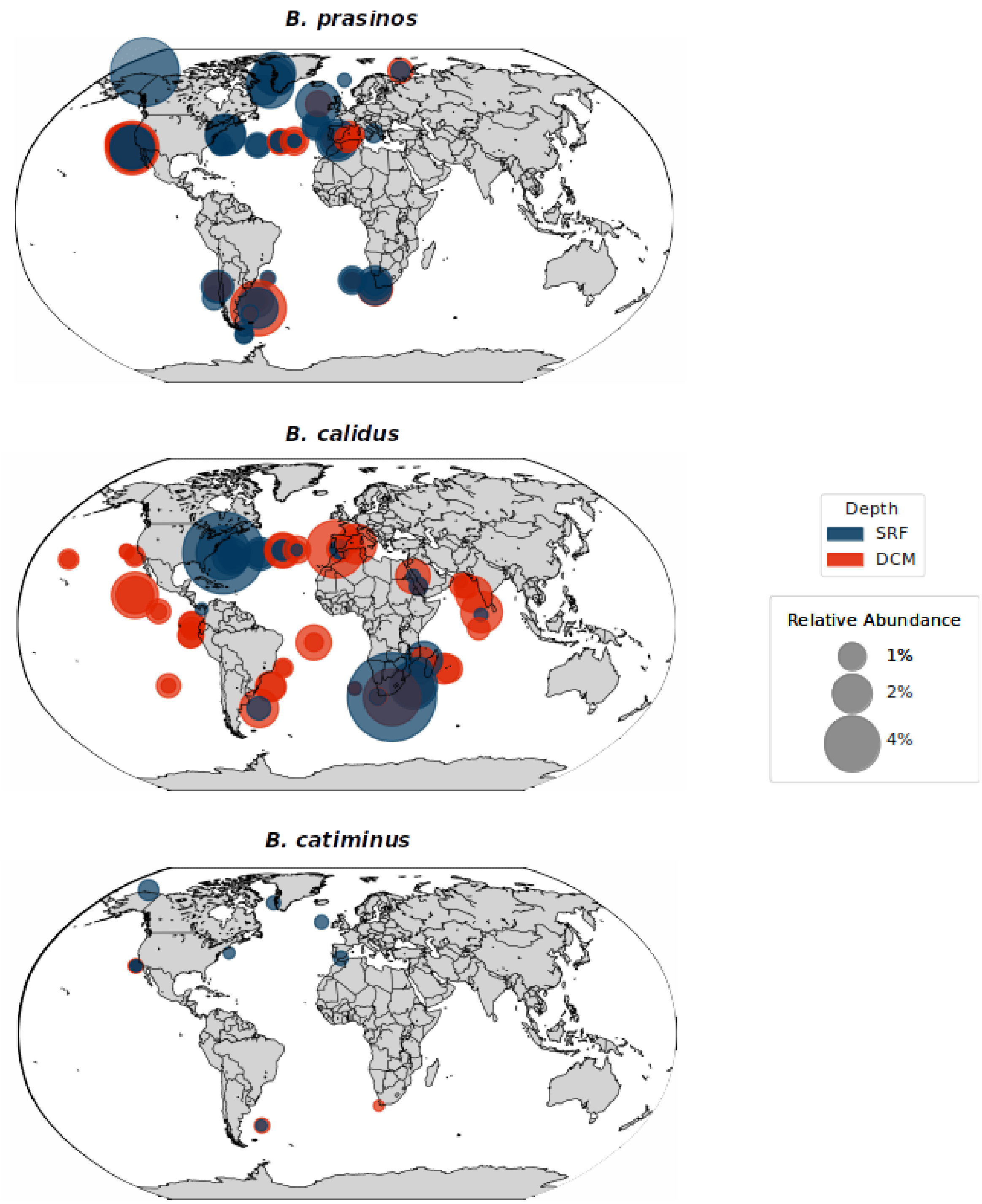


**Fig S4: Biogeography of the *Bathycoccus* genus.** Geographical distribution of *Bathycoccus* species based on read mapping from TARA ocean metagenomic datasets. For each species, only stations with horizontal genome coverage higher than 10% and 4 reads of minimal depth are displayed. Relative abundance corresponds to the percentage of reads from the dataset mapped to the genome.

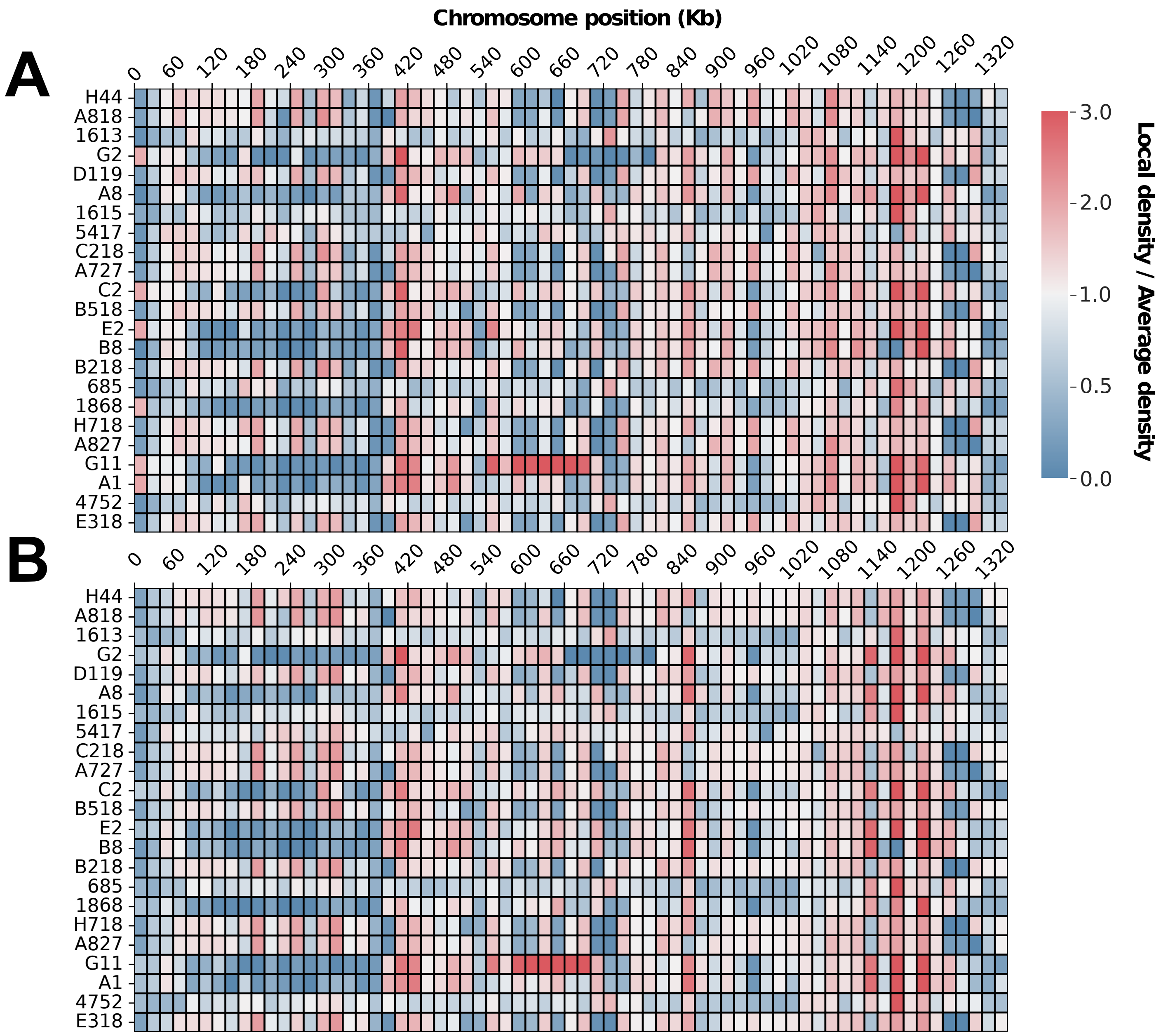


**Fig S5: Genomic diversity of Chromosome 1 in *Bathycoccus prasinos*.** (A) Divergences between local SNP density and genome wide SNP density and (B) between local SV density and genome wide SV density along a 20 kb sliding window. Variant calling was computed against strain RCC4222.

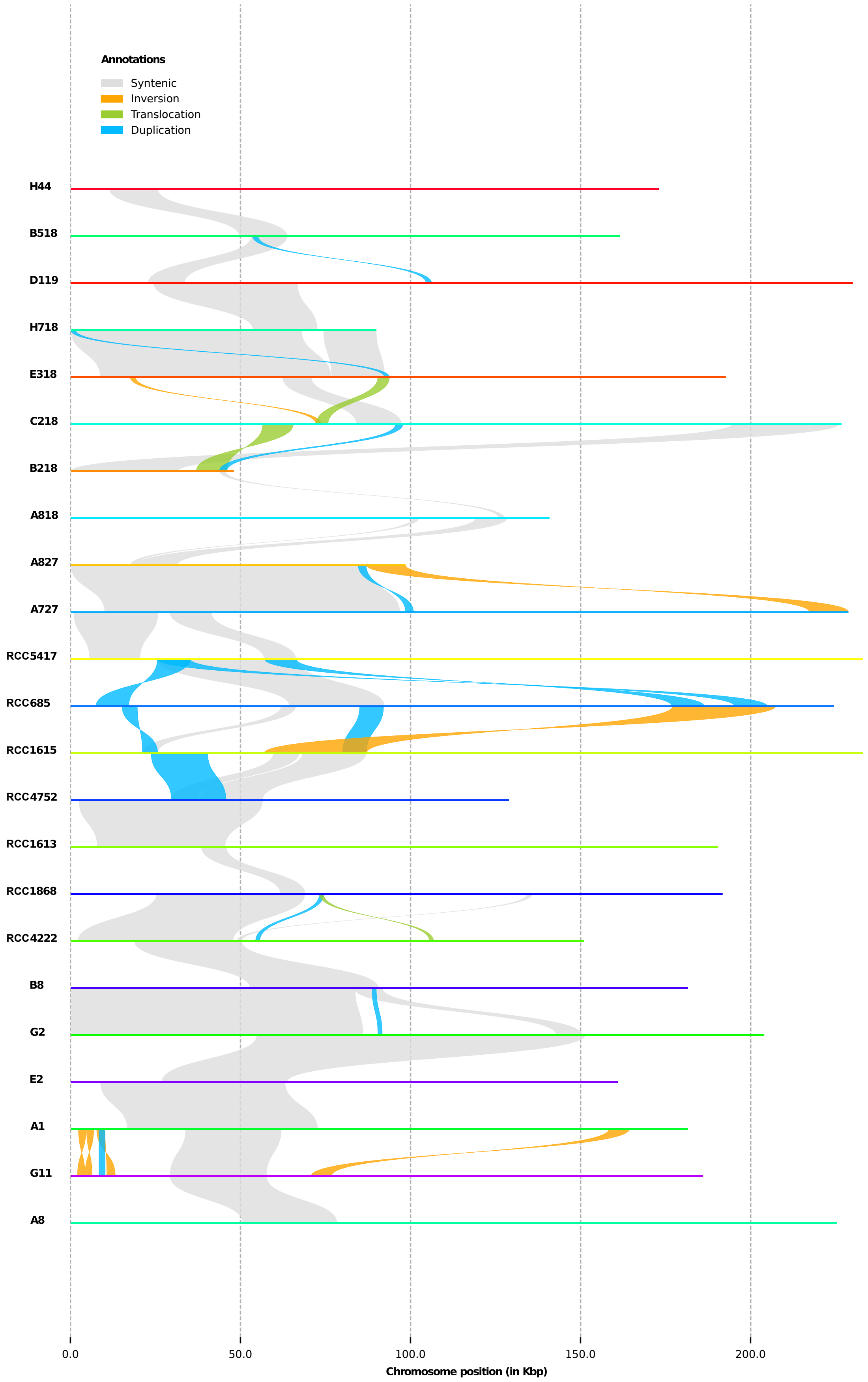


**Fig S6: Chromosome 19 pairwise alignments between *B. prasinos* strains.** Chromosome 19 alignment and structural variation visualization between *B. prasinos* assemblies. Strain order follows phylogenetic order shown in Fig. 4. Strain C2 was not displayable due to the lack of shared sequences with phylogenetically close strains. Only structural variations longer than 500 bp are displayed.
